# Bayesian inference of phylogeny is robust to substitution model over-parameterization

**DOI:** 10.1101/2022.02.17.480861

**Authors:** Luiza Guimarães Fabreti, Sebastian Höhna

## Abstract

Model selection aims to choose the most adequate model for the statistical analysis at hand. The model must be complex enough to capture the complexity of the data but should be simple enough to not overfit. In phylogenetics, the most common model selection scenario concerns selecting an appropriate substitution and partition model for sequence evolution to infer a phylogenetic tree. Here we explored the impact of substitution model over-parameterization in a Bayesian statistical framework. We performed simulations under the simplest substitution model, the Jukes-Cantor model, and compare posterior estimates of phylogenetic tree topologies and tree length under the true model to the most complex model, the GTR+Γ+I substitution model, including over-splitting the data into additional subsets (*i.e*., applying partitioned models). We explored four choices of prior distributions: the default substitution model priors of `MrBayes, BEAST2` and `RevBayes` and a newly devised prior choice (`Tame`). Our results show that Bayesian inference of phylogeny is robust to substitution model over-parameterization but only under our new prior settings. All three default priors introduced biases for the estimated tree length. We conclude that substitution and partition model selection are superfluous steps in Bayesian phylogenetic inference pipelines if well behaved prior distributions are applied.

## 1 Introduction

At the heart of all model-based phylogenetic inferences lies the substitution model. The substitution model defines the rate of substitutions between any pair of states (*e.g*., between nucleotides) and thus the probabilities of substitutions given a branch length. Many different substitution models have been suggested over time, *e.g*., the Jukes-Cantor (JC) model (Jukes and Cantor 1969), the Kimura two parameter (K2P) model (Kimura 1980), the Kimura three parameter (K3P) model (Kimura 1981), the Felsenstein (F81) model (Felsenstein 1981), the Hasegawa-Kishino-Yano (HKY85) model (Hasegawa et al. 1985), and the general time reversible (GTR) model (Tavaré 1986). Additionally, phylogenetic substitution models often incorporate different rates among sites (the +Γ model, Yang 1994) and/or a proportion of invariable site (the +I model, Adachi and Hasegawa 1995; Gu et al. 1995). With this diversity of substitution models, a researcher is left with the daunting task to choose the “best” substitution model for the specific dataset at hand.

An *under-parameterization* (*i.e*., oversimplified) substitution model can bias phylogeny estimation (Sullivan and Joyce 2005). This problem was demonstrated in several applications, *e.g*., Leitner et al. (1997), Sullivan and Swofford (1997), Cunningham et al. (1998), Kelsey et al. (1999). These studies concluded that phylogenetic inference with different substitution models can result in a significant difference of the tree topology and/or branch lengths, with the more complex models performing better in all cases. The observed biases introduced due to under-parameterized substitution models has led to the development of methods for substitution model selection (Posada and Crandall 1998; Huelsenbeck et al. 2004; Posada 2008; Darriba et al. 2012; Kalyaanamoorthy et al. 2017). It has become common practice to select the best fitting substitution model before estimating a phylogenetic tree. For example, in 2014, the paper describing the software `ModelTest` (Posada and Crandall 1998) was among the the 100 most cited papers of all time (Van Noorden et al. 2014).

There are several problems with the current approach of substitution model selection in phylogenetics pipelines. First, the current approach is circular because the model selection step (*e.g*., in `ModelTest` and its successors, Posada and Crandall 1998; Posada 2008; Darriba et al. 2012) requires a phylogeny (Bouckaert and Drummond 2017). This phylogeny is often estimated using fast but less accurate models and methods (*e.g*., using Neighbor-Joining, Posada and Crandall 2001). Using the wrong phylogeny could lead to biased model selection. Second, the current approach does not incorporate uncertainty in the estimated phylogeny, branch lengths and substitution model parameters (Nylander et al. 2004; Bouckaert and Drummond 2017). The crucial issue with substitution model selection is that considerable shortcuts are taken because the analysis using a single substitution model can take weeks to months. Performing full inferences and model selection, *e.g*., computing Bayes factors for each substitution model (Höhna et al. 2017), increases the computation time by several factors even using parallel computations (Höhna et al. 2021). Full substitution model selection is infeasible and (almost) never applied because of this computational demand.

Is substitution model selection in Bayesian phylogenetic inference a necessary step, or could simply the most complex, *e.g*., the GTR+Γ+I substitution model, be used? It has been shown that an *under-parameterized* substitution model can bias phylogeny estimation (see Sullivan and Joyce 2005, for a review) but does an *over-parameterized* (*i.e*., too complex) substitution model also biases phylogenetic inference? Surprisingly, this question has received rather little attention and only two studies have partially addressed this question (Huelsenbeck and Rannala 2004; Lemmon and Moriarty 2004). First, Huelsenbeck and Rannala (2004) studied the posterior probabilities of bipartitions under simple and complex substitution models. They concluded that Bayesian inference is more sensitive to under-parameterization than to over-parameterization with regard to tree topology. Second, Lemmon and Moriarty (2004) specifically studied the impact of model misspecification on Bayesian inference of phylogeny. They show that model over-parameterization does not bias bipartition posterior probabilities and has little to no effect on branch lengths. The observed effect on branch lengths was a decrease in precision. Therefore, Lemmon and Moriarty (2004) conclude with a warning to not assume the most complex model because of the “imprecision that may result from over-parameterization”. Thus, substitution model selection remains common practice in phylogenetic inference.

Furthermore, in most phylogenetic analyses, the sequence alignment is divided into several data subsets, *e.g*., a multi-locus dataset divided by gene or codon position. Each data subsets can either receive its own substitution model or share the substitution model with another data subset (so-called partition models, Nylander et al. (2004)). The partition model accommodates process heterogeneity along molecular data and improper data partitioning and application of under-parameterized substitution model can bias phylogenetic inferences (Brown and Lemmon 2007). The number of possible partition models to choose from is significantly larger than the number of available substitutions models. If selecting the best substitution model for a single locus is already burdensome and computationally extremely demanding, then selecting the best partition model is clearly infeasible without even more drastic shortcuts. Nevertheless, several methods have been developed for partition model selection (*e.g*., Lanfear et al. 2012, 2017) and are commonly applied in phylogenetic inference pipelines. Until today, there has been no study to evaluate if over-parameterization, *i.e*., assuming a separate GTR+Γ+I substitution model per data subset, biases phylogenetic inference.

In this study, we will investigate the effect of model over-parameterization on Bayesian phylogenetic analysis. Specifically, we focus on the question if substitution model selection is a necessary step for Bayesian phylogenetic inference. Can we simply use the most complex substitution model and partition model and thus avoid the danger of under-parametrized models? Here, we explore this question using simulations. We simulated data under the simplest model and inferred the phylogenies under (a) the true model, (b) an over-parameterized substitution model, and (c) an over-parameterized partition model. Moreover, we tested different choices of prior distributions for the over-parametrized substitution model. The advantage of using simulations over previous studies using empirical data (*e.g*., Abadi et al. 2019) is that we know the true parameters (*i.e*., the true phylogeny and branch lengths) and the true model. Therefore, we are able to assess if over-parametrization biases our results or leads to less precise estimates (*i.e*., higher uncertainty and larger credible intervals). We assessed biases by comparing bipartition posterior probabilities and tree lengths between analyses under the true substitution model and an over-parametrized model.

## 2 Methods

We performed a simulation study to understand how the estimated tree topology and tree length are affected under an over-parameterized substitution model (within the GTR family of nested substitution models). Our approach is depicted in Figure 1. The data sets were simulated under the simplest substitution model, the Jukes-Cantor substitution model (JC). We over-parameterized the substitution model by using the most parameter rich commonly used substitution model, the GTR+Γ+I substitution model, to perform the phylogenetic inference. The simulation with the simplest substitution model in combination with the inference with the most complex comprise the most extreme case of over-parameterization of common substitution models, and therefore the most conservative scenario to evaluate the effects of substitution model over-parameterization. If we find no impact of substitution model over-parameterization for this extreme scenario, then there is no impact of substitution model over-parameterization for less extreme cases.

**Figure 1:**
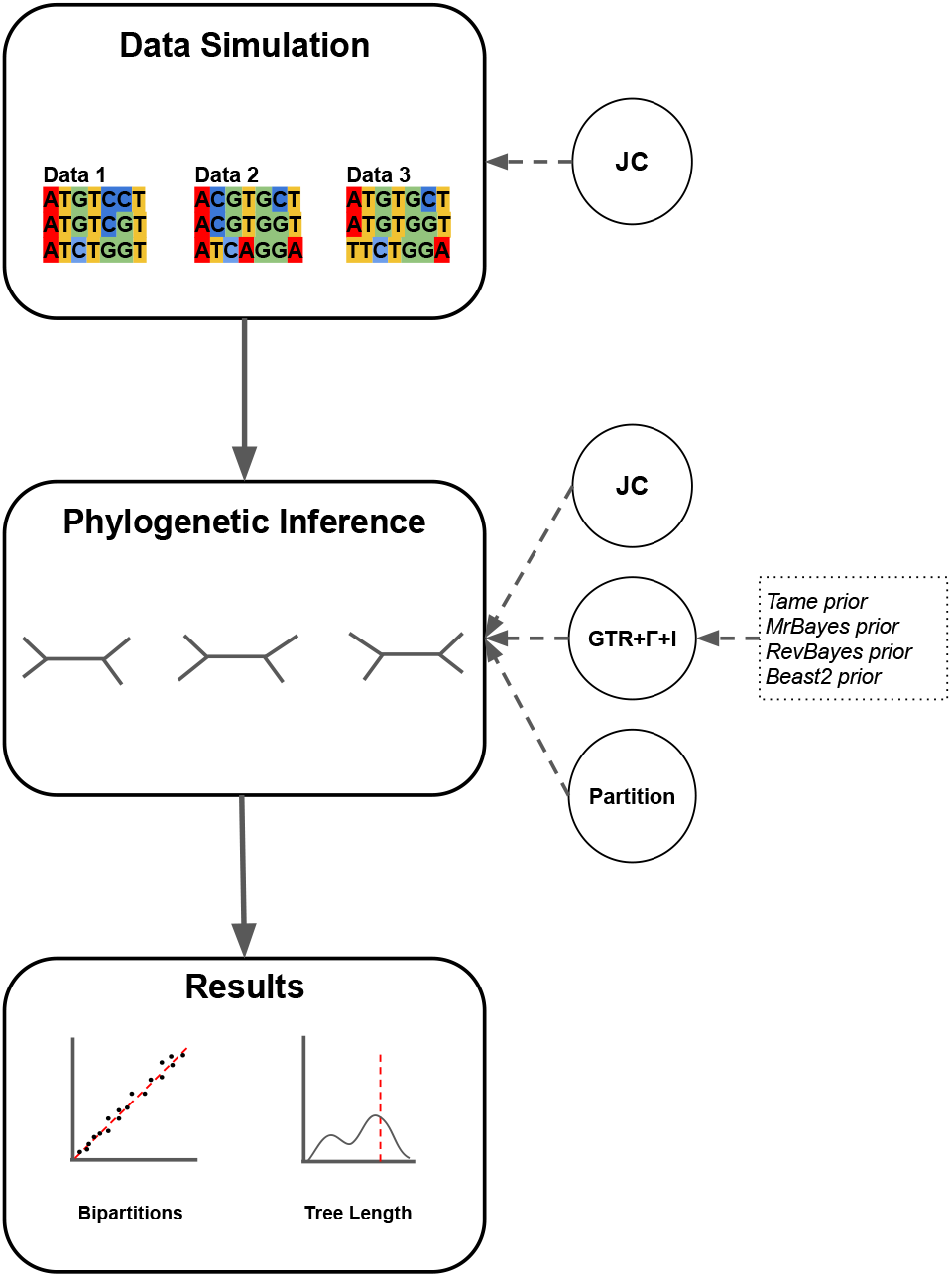
Summary of the simulation and analyses used in this study. The first step contains the simulation of data matrices under the Jukes-Cantor (JC) substitution model. The next step contains the phylogenetic inference under the different settings (GTR+Γ+I with four different prior settings and the partition model). The comparisons between the true and over-parameterized models are summarized in the results.

We varied the prior probability distributions for the over-parameterized model according to the default settings of three commonly used Bayesian phylogenetic software (`MrBayes, BEAST2` and `RevBayes`). These default prior distributions for the Bayesian phylogenetic software were extracted for: a) `MrBayes` v3.2 (Ronquist et al. 2012); b) the `RevBayes` protocol described in Höhna et al. (2017); c) `BEAST2` using BEAUTi (Bouckaert et al. 2019). The difference in default prior distributions among these popular software packages reflects the uncertainty about good prior choices in the field, and our choice of these three example does not represent a favor for or against any of these choices. We included another prior distribution (called “`Tame`” in this manuscript). Additionally, we performed inference under an over-partitioned scheme for the novel prior setting where each data subset received its own separate GTR+Γ+I substitution model. Next, we compared posterior probabilities of bipartitions and tree length credible intervals between the models used for inference. The following sections explain in more detail the methods used in this study.

### 2.1 Simulation Settings for the Datasets

We simulated data matrices by first drawing all parameters from their prior distribution. The substitution model was set to the JC model of sequence evolution (Jukes and Cantor 1969). The JC substitution model is the simplest model within the GTR family of nested models and has no free substitution model parameters (all base frequencies are fixed to 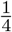 and all relative exchangeability rates are fixed to 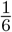). Therefore, the prior distributions for the simulations were the tree topology prior and the branch lengths prior. We assumed a uniform distribution on tree topology, *i.e*., each topology had equal prior probability. We chose two different tree sizes, 16 and 64 taxa, to explore the impact of tree size. The prior distribution for the branch lengths was an exponential distribution with either a mean of 0.1 or 0.02, which impacts the total number of substitutions expected along the phylogeny and therefore the informativeness of the data sets (Figure S1). We defined the number of sites to be 100 or 1000 to explore the impact of the amount of data. In summary, we simulated data sets under all possible combinations of mean branch length, number of taxa and number of sites, yielding eight different simulation scenarios which are displayed in Table 1. We simulated 1000 replicates for the data set with 16 taxa and 500 replicates for the data sets with 64 taxa. The simulations were performed using RevBayes (Höhna et al. 2016) and all scripts are available at https://github.com/lfabreti/SM-over-parameterization.

**Table 1:** The different settings for the simulation of data matrices under the Jukes-Cantor substitution model. The first column shows the index of the simulation setting. The second column is the number of taxa, followed by the number of sites and the mean branch length. The gray cells highlight the changes in the settings for each simulation setting.

| Simulation setting | Number of taxa | Number of sites | Mean branch length |
| --- | --- | --- | --- |
| 1 | 16 | 100 | 0.1 |
| 2 | 16 | 100 | 0.02 |
| 3 | 16 | 1000 | 0.1 |
| 4 | 16 | 1000 | 0.02 |
| 5 | 64 | 100 | 0.1 |
| 6 | 64 | 100 | 0.02 |
| 7 | 64 | 1000 | 0.1 |
| 8 | 64 | 1000 | 0.02 |

### 2.2 Prior Distributions on the Substitution Model Parameters

In Bayesian inference, the prior probability distribution defines the researcher’s belief about the parameter quantity before the data are taken into account. All parameters and hyper-parameters from a model need to be assigned to a prior probability distribution. The best approach to define a prior probability distribution for a given parameter is an open debate in Bayesian inference (Lindley 1961; Morris et al. 2015; Gelman and Hennig 2017; Lemoine 2019; Banner et al. 2020). In some situations in phylogenetics, the researcher has reliable information to make strong assumptions about the prior distribution for a given parameter. For example, informative prior distribution are commonly used in divergence time estimation using node calibrations where fossil information is used to define minimum and maximum bounds on the age of a given node (Parham et al. 2012; Warnock et al. 2012). If no reliable information about parameter values is available, then the prior should be designed to have little effect on the estimated parameters (Zwickl and Holder 2004; Alfaro and Holder 2006). Next, we will discuss the default priors for the GTR+Γ+I model adopted by three commonly used Bayesian phylogenetic software (`MrBayes, BEAST2` and `RevBayes`) and our proposed prior scheme (`Tame`).

The GTR model has four equilibrium base frequencies and six rates of changes between bases (exchange-ability rates). The among site rate variation model (ASRV) and the invariant sites model have each one parameter, the shape parameter *α* and the probability of a site being invariant, respectively. All default prior distributions assign equal probabilities for all four base frequencies (Table 2). The main differences in the prior schemes assessed here lie in the exchangeability rates and the *α* parameter of the among site rate variation model. To aid grasping the impact of the different prior distribution choices, we provide figures depicting the behavior of each induced parameter given the prior settings (Figures 2 and 3).

**Table 2:** Description of the (default) prior settings for the commonly used phylogenetic tools and the proposed prior. The first column displays the name of the prior setting. The following columns are the parameters for the GTR+Γ+I model.

| Prior | Branch length | Equilibrium base frequencies | Exchangeability rates | Alpha parameter | Proportion of invariant sites |
| --- | --- | --- | --- | --- | --- |
| **Tame** | Exponential ($\lambda=10$) | Dirichlet (1,1,1,1) | Dirichlet (1,1,1,1,1,1) | Uniform (0, $1 \times 10^8$) | Beta ($\alpha=1, \beta=1$) |
| **MrBayes** | Exponential ($\lambda=10$) | Dirichlet (1,1,1,1) | Dirichlet (1,1,1,1,1,1) | Uniform (0,200) | Uniform (0,1) |
| **RevBayes** | Exponential ($\lambda=10$) | Dirichlet (1,1,1,1) | Dirichlet (1,1,1,1,1,1) | Lognormal ($\mu=\ln(1.5)$, $\sigma=0.587$) | Beta ($\alpha=1, \beta=1$) |
| **BEAST2** | Exponential ($\lambda=10$) | Uniform (0,1) | Gamma (k=0.05, $\theta=10$ or 20) | Exponential ($\lambda=1$) | Uniform (0,1) |

**Figure 2:**
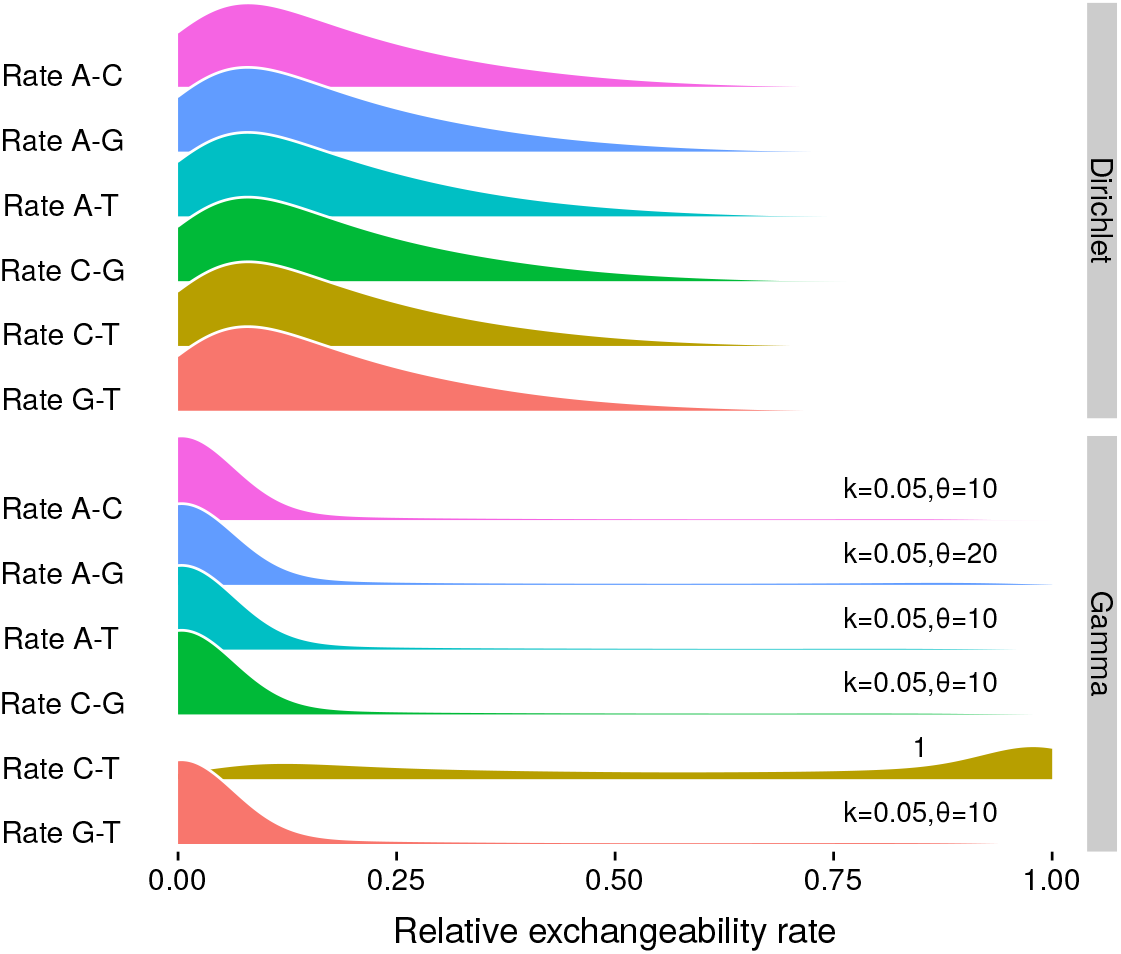
The prior distribution of the exchangeability rates. Here we show the two commonly used prior distributions on exchangeability rates; the Dirichlet distribution and the gamma distribution. The gamma distribution has a shape 0.05 and scale parameter 10 for the rates A↔C, A↔T, C↔G and G↔T; the rate A↔G has a different scale of 20; the rate C↔T is set to 1 for normalizing the rates. These densities are produced by simulating 1 × 10^6^ samples from the corresponding distributions.

**Figure 3:**
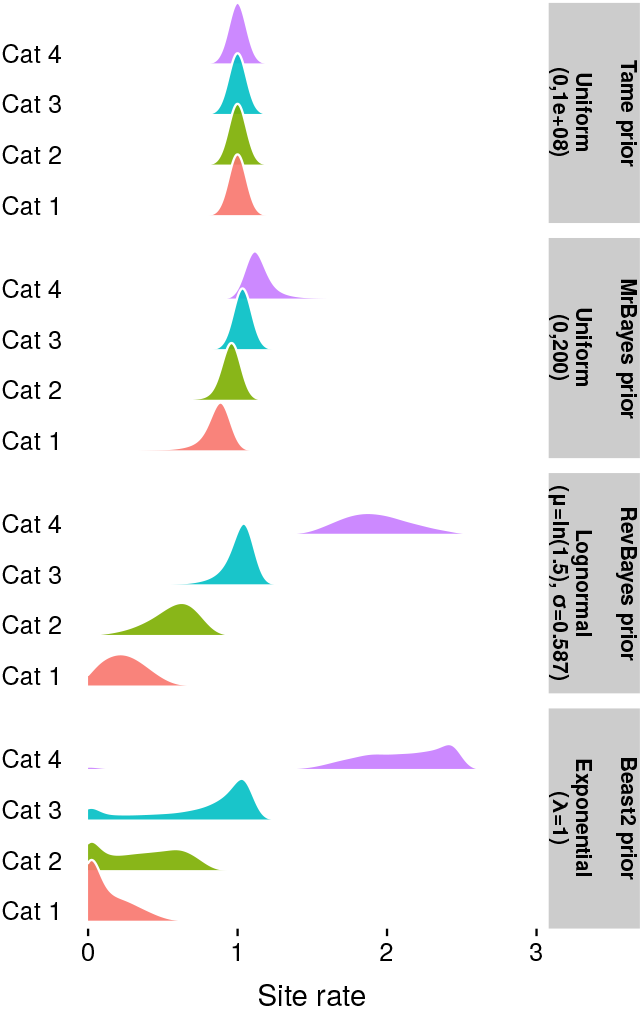
The induced prior distributions of the four rate categories from the among site rate variation model for the different *α* priors. The first panel draws *α* from a uniform distribution from 0 to 1 10^8^ and it is the proposed prior distribution in this study. The second panel uses a uniform form 0 to 200 as the prior on *α*, which is implemented in `MrBayes`. The third panel shows the default behavior for `RevBayes`, where *α* is drawn from a lognormal distribution with *μ*=ln(1.5) and *σ*=0.587405. The last panel shows the behavior of the four gamma categories when the prior on *α* is set to an Exponential distribution with *λ*=1, as it is done in `BEAST2`.

We compared the profile of two different prior distributions on the exchangeability rates, a flat Dirichlet(1,1,1,1,1,1) distribution and a gamma(k=0.05, *θ*=10 or 20) distribution. The prior schemes for the `Tame, MrBayes` and `RevBayes` use the former, while `BEAST2` assumes the latter. Note that the prior settings `Tame, MrBayes` and `RevBayes` require that the sum of all rates equals to 1.0 while `BEAST2` rescales the rates internally and fixes the rate between C↔T to 1.0. Figure 2 shows the distribution for the six exchangeability rates for each prior assumption. The Dirichlet prior distribution results in an equal distribution with a mean of ≈ 0.16 for all six rates, which is expected since the concentration parameter is the same for all categories. The gamma prior distribution used in `BEAST2` yields four equal distributions with a mean of *≈* 0.07 for the rates A↔C, A↔T, C↔G and G↔T. The induced prior distribution for the rate A↔G is slightly larger due to the scale parameter *θ*=20, which results in a mean of *≈* 0.1. We observed the largest difference for the relative rate C↔T with a mean of *≈* 0.6 and the values are concentrated on higher rates. Note that the rate C↔T is originally set to 1.0 and used to normalize the exchangeability rates. All other unscaled exchangeability rates have a prior mean of 1.0 (for A↔G) and 0.5 (for all other rates) but an induced relative mean which is clearly different from 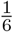.

The prior distributions for the shape parameter, *α*, of the gamma distribution used to model among site rate variation, differs considerably among standard Bayesian phylogenetic inference software (Table 2). `MrBayes` v3.2 uses a uniform(0,200) distribution as a prior distribution for *α*, which has a mean of 100. Note that more recent versions of `MrBayes`, starting with version 3.2.2, assume an exponential prior distribution. The RevBayes protocol establishes a biologically motivated prior distribution as a lognormal distribution with median 1.5 and standard deviation 0.587405 (Höhna et al. 2017). This prior distribution was designed to specify a 95% prior distribution which ranges from a 3-fold to 100-fold difference between the lowest and highest rate categories (Höhna et al. 2017). `BEAST2` uses by default an exponential(*λ*=1) prior distribution for *α* with a mean of 1.0.

The induced prior distributions of the four rate categories for each prior distribution on *α* are shown in Figure 3. One might expect that small values for *α*, or even converging towards 0.0, result in no rate variation. However, the contrary is true. Smaller values of *α* result in more distinct distributions for the four discretized gamma quartiles, which translates into more rate variation across sites. Both the `RevBayes` and `BEAST2` prior settings assign more prior probability to smaller values for *α* resulting in larger variation between the rate categories. The `MrBayes` prior setting results in more uniformity among the categories, but still expecting some rate variation *a priori*. We propose a prior that results in less *a priori* expected rate variation between the rate categories, namely a uniform(0, 1 *×* 10^8^) distribution (Figure 3 upper panel). This exploration shows the usefulness to examine the induced prior distribution of the model parameters.

### 2.3 Phylogenetic Inference Settings

Phylogenetic inference was performed in a Bayesian Markov chain Monte Carlo (MCMC) framework implemented in `RevBayes` and MCMC convergence was assessed using the `R` (R Core Team 2020) package `Convenience` (Fabreti and Höhna 2022). Each simulated dataset was analyzed under two substitution models: either JC or GTR+Γ+I. The JC substitution model represents the true model whereas the GTR+Γ+I represents the over-specified substitution model. We applied four different prior schemes to the GTR+Γ+I, as described in Table 2. The total number of replicated inferences per simulated condition was 1000 for the data with 16 taxa and 500 for the data with 64 taxa. Additionally, we used a partition model, for the data sets with 1000 sites, with two equal-sized data subsets evolving independently under a GTR+Γ+I with the `Tame` prior scheme. For these partition models we only used the simulated data with 1000 because splitting 100 sites results into unrealistically small data subsets. In this case, we performed 500 inference replicates for the data with 16 taxa and 300 for the data with 64 taxa. We used two replicated MCMC runs for each inference and samples were taken at every 20^th^ iteration. The criteria for convergence assessment were the default from Convenience as described in Fabreti and Höhna (2022). This strict convergence assessment turned out very useful as several runs showed outlier behavior which could be attributed to non-convergence.

The total number of MCMC iterations varied based on the convergence status. We started with 100,000 iterations and increased this value for analyses that did not converge. The maximum number of iterations used was 400,000. The moves during the MCMC followed the scheme on Table 3. The moves on the tree parameter varied according to the size of the dataset. For a dataset with 16 taxa, each iteration had 76 moves, whereas a dataset with 64 taxa had 244 moves per iteration. The inference with the `BEAST2` prior setting yielded a different move scheme for the exchangeability rates because each parameter had its independent prior distribution instead of a compound prior distribution. In this case, each of the estimated rates (A↔C, A↔G, A↔T, C↔G and G↔T) was assigned with a scale move with weight two.

**Table 3:** Description of the move settings during the MCMC. Each row corresponds to a model parameter, the second column shows the move applied to the parameter and the third column shows the weight for the moves. The tree topology moves were the Nearest Neighbor Interchange (NNI) and Subtree Pruning and Re-Grafting (SPR).

| Parameter | Move | Weight |
| --- | --- | --- |
| Tree | NNI (Nearest Neighbor Interchange) | # taxa |
| | SPR ( Subtree Pruning and Re-Grafting) | $\frac{\# \text{ taxa}}{2}$ |
| Branch length | Branch length scale | # branches |
| Equilibrium frequencies | Beta simplex | 4.0 |
|  | Dirichlet simplex | 2.0 |
| Exchangeability rates | Beta simplex | 6.0 |
|  | Dirichlet simplex | 3.0 |
| Exchangeability rates (BEAST2) | 5 scale moves | 2.0 |
| Alpha | Scale | 4.0 |
| Proportion of invariant | Beta probability | 4.0 |

### 2.4 Evaluation of bias and uncertainty in estimated parameters

The two key parameters of interest for most phylogenetic studies are the tree topology and the branch lengths. Therefore, we focused our evaluation of the effect of over-parameterization on these two parameters. The tree topology was translated into its bipartitions, which are subsets of the full tree. We analyzed two outcomes of the MCMC output: (1) the posterior probability of bipartitions and (2) the 95% credible interval for the tree length. These estimates were compared between the inference under the true model (JC) and the over-parameterized models (GTR+Γ+I). If over-parameterization is not problematic, then the estimates under the GTR+Γ+I model will not deviate from the estimates under the JC substitution model.

The posterior probability of any given bipartition was compared to whether the bipartition was true, *i.e*., present in the true tree. Thus, we obtained the frequency of a bipartition being true given its posterior probability. For a more stable computation of the frequencies, we binned bipartition into 20 equal-sized bins for posterior probabilities between 0.0 and 1.0, *e.g*., the first bin for bipartitions with posterior probabilities 0.0 ≤ *PP* < 0.05. The expected behavior is that the overall frequency over all replicates of a bipartition being true is equal to the posterior probability (Huelsenbeck and Rannala 2004). For example, if we observe thousands of bipartitions with probability *ξ*, *e.g*., 0.2, then we expect *ξ* percent of these bipartitions to be true, *e.g*., 20% of the bipartitions. We evaluated the behavior of the posterior probabilities vs. frequency of being true for all five inference scenarios.

The 95% credible interval was used as a measure of the precision of the estimated tree length. Larger credible intervals imply more variance in the estimated tree length. The reference for the size of the credible interval was the inference under the true model, *i.e*., the model used for simulating the data (JC). We compared the size of the credible interval for the tree length between the analysis under JC and GTR+Γ+I, with the four different prior schemes. Additionally, we explored further the posterior probability distributions of the tree length across the inference settings by examining one example dataset for each simulated scenario. If over-parameterization is not a problem, then the estimated 95% credible interval between the inferences under the JC substitution model (the true model) and the GTR+Γ+I substitution model (the over-parameterized model) should match and the estimates should be seen on the 1:1 line. Conversely, we would expect that if over-parameterization adds uncertainty in our estimates, then we should obtain larger 95% credible intervals for the analyses under the GTR+Γ+I substitution model.

## 3 Results

### 3.1 Accuracy of Posterior Probability of Bipartitions

We observed no major deviation from the expected behavior under all conditions (Figure 4, the expected behavior was that the bipartitions fall on the 1:1 line). The minor variation observed in Figure 4 is due to the intrinsic randomness of the bipartition posterior probability estimates obtained from the MCMC samples and is also observed for the inference under the true model (JC). Adding more replicates would improve the fit to the diagonal line, but the underlying behavior is already observable with the current amount of replicates, which took several months on our local High Performance Cluster to complete. We note that estimated bipartition probabilities for the simulations with 64 taxa are more accurate, which is due to the larger number of bipartitions in each dataset.

**Figure 4:**
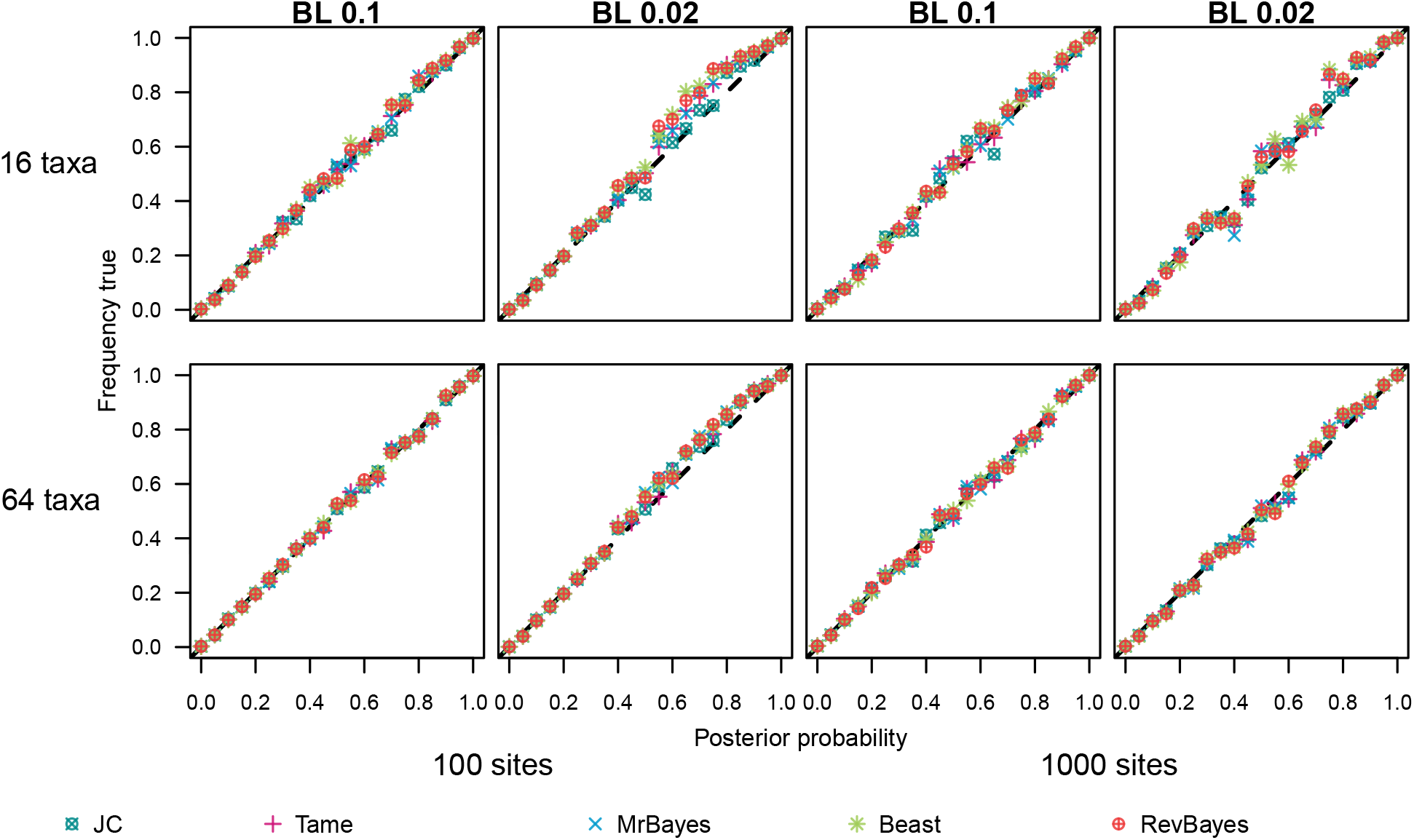
The relationship between posterior probability of bipartitions and the bipartition was correct. The expected behavior is that, on average, the posterior probability and the frequency how often a bipartition with this posterior probability is indeed correct are exactly correlated following the 1:1 line. The different panels show each of the 8 settings in which the data were simulated. The first row corresponds to the simulated trees with 16 taxa, while the second row corresponds to the simulated trees with 64 taxa. The mean branch lengths (BL) for the data sets are on top of each column. The two first columns display the data sets with 100 sites, the two other columns show the data sets with 1000 taxa. The symbols represent the different models and prior settings used for the inference (Table 2). All models, including the over-parameterized models, produced statistically consistent estimates of the bipartition posterior probabilities.

Our results agree with previous observations by Huelsenbeck and Rannala (2004) and Lemmon and Moriarty (2004). These two previous studies also showed that substitution model over-parametrization has no effect on estimated bipartition posterior probabilities. Our study clearly corroborates this finding; even a severe over-parameterization of the substitution model does not impact estimated bipartition posterior probabilities. Furthermore, we observed no difference in the estimated bipartition posterior probabilities based on the prior distribution setting used. We also did not observe any impact of the alignment length, number of taxa and branch length prior on the accuracy of the estimated bipartition posterior probabilities. This indicates that, at least for our simulations, the choice among common default prior distributions on the substitution model parameters does not impact the accuracy of bipartition posterior probabilities.

Similarly, we observed no biases in the posterior probabilities of bipartitions (Figure 5) for the inference under the partition model. Figure 5 shows the same variation around the diagonal line as Figure 4 due to the intrinsic stochasticity of the MCMC samples and is also observed for the inference under the true model (JC). This effect is larger for the simulated data sets with 16 taxa, because these simulations have less bipartitions in each replicate. These results for the over-splitted partition model are not surprising as we have seen in Figure 4 that even very small datasets of only 100 sites are not impacted by substitution model over-parameterization. Therefore, it is logical that over-splitting and over-parameterization of partition models is not a problem if the size of each data subset is not too small. We conclude that over-parameterization of the substitution and partition models does not affect the posterior probability of bipartitions, and therefore, the tree topology.

**Figure 5:**
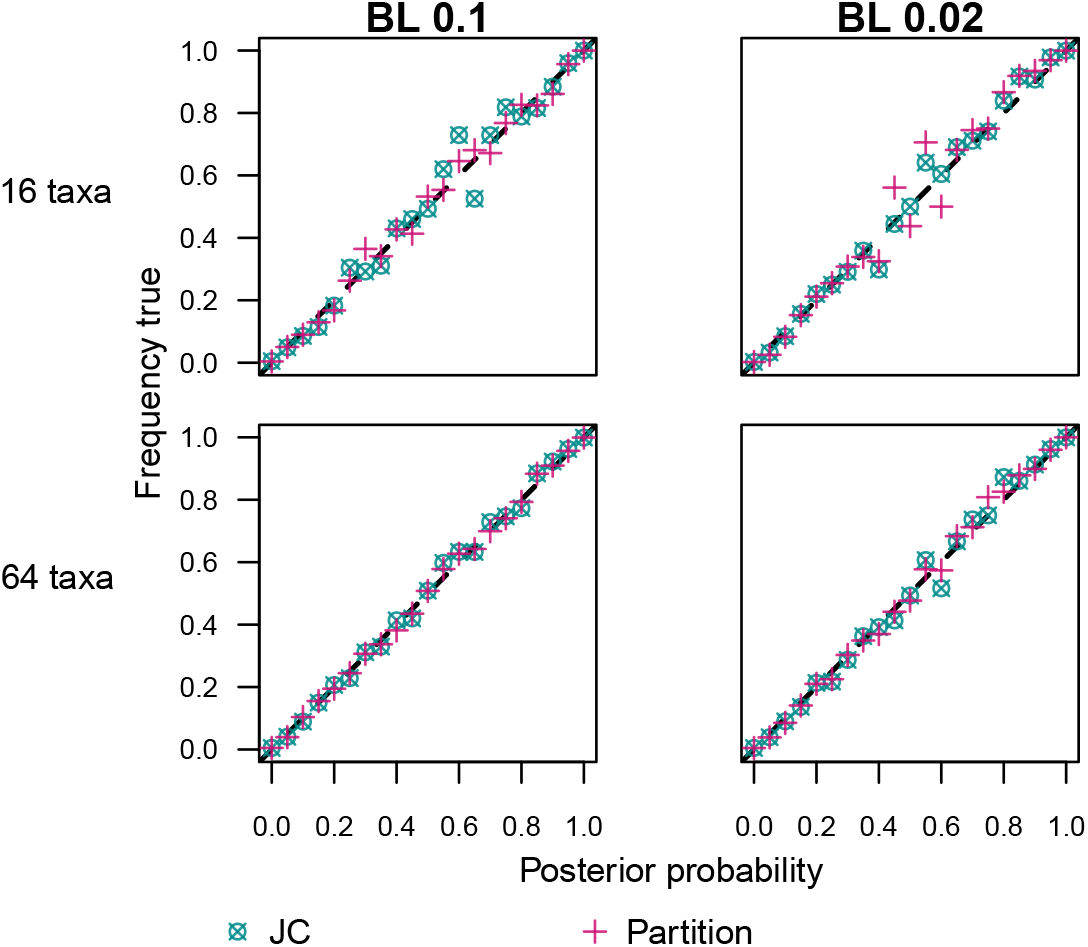
The relationship between posterior probability of bipartitions and the bipartition being correct under the partition model. The expected behavior is that the posterior probabilities of the bipartitions and the frequency that a bipartition with this posterior probability was correct follows the 1:1 line. The different panels show each of the 4 settings in which the data was simulated. The first row corresponds to the simulated trees with 16 taxa, while the second row corresponds to the simulated trees with 64 taxa. The mean branch lengths (BL) for the data sets are on top of each column. For the over-partitioned model, we only used data sets with 1000 sites. The symbols represent the models for the inference, JC or the over-partitioned GTR+Γ+I model (using the `Tame` prior settings for both data subsets).

### 3.2 Accuracy of Estimated Tree Length

We observed an impact of substitution model over-parametrization on the width of the credible interval of the tree length (Figure 6). The 95% credible interval of the tree length was very similar between JC and GTR+Γ+I for all simulated matrices with 1000 sites (Figure 6). However, with less data (100 sites), the choice of prior distribution had an observable impact on the estimated tree length and the resulting 95% credible interval (Figures S2-S5). Only our newly proposed `Tame` prior settings produced unbiased estimates of the 95% credible interval.

**Figure 6:**
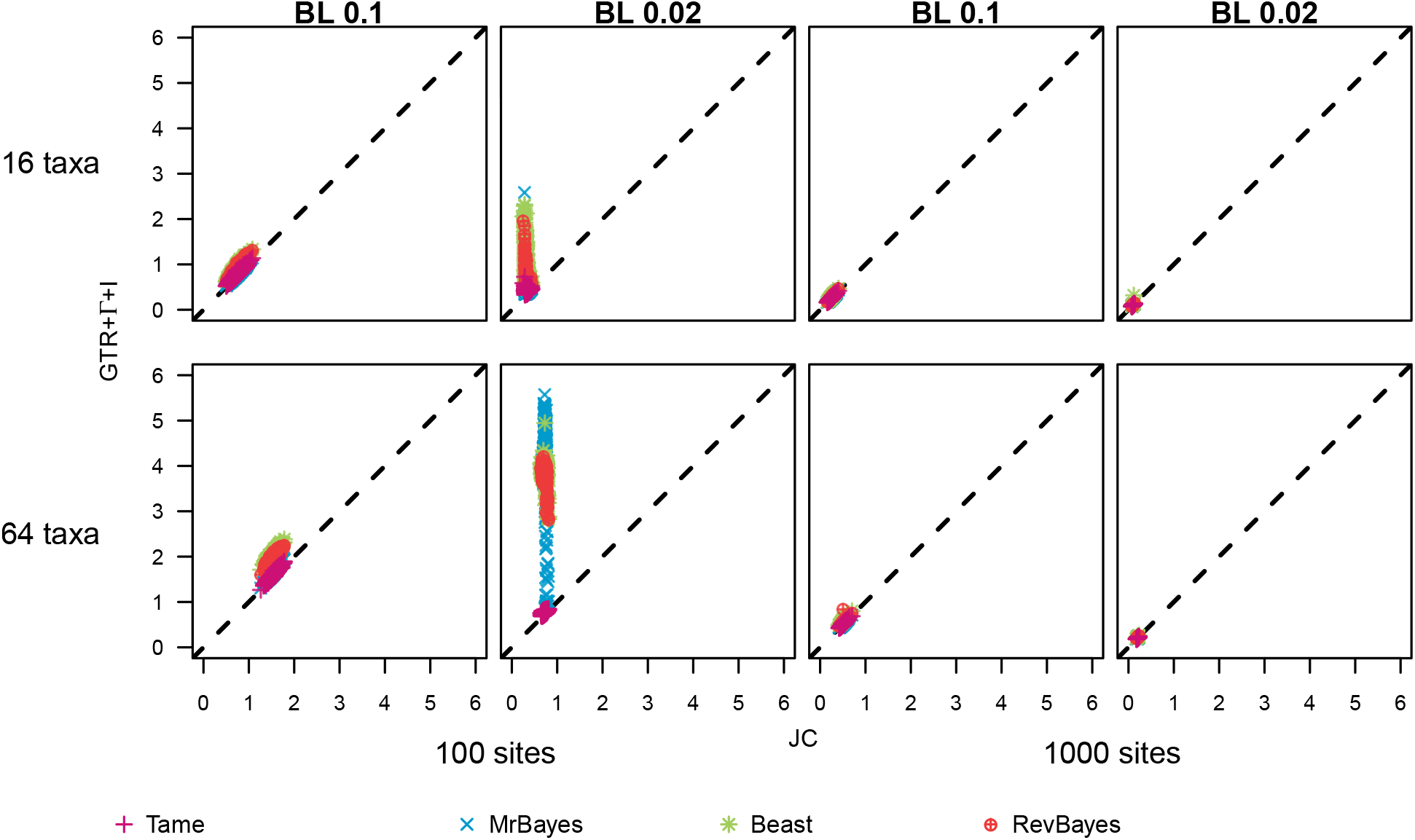
The width of the 95% credible interval for the tree length for the inference with Jukes-Cantor (JC) against the inference with GTR+Γ+I. On the x-axis we show the estimated 95% credible interval size for the JC substitution model (true model). On the y-axis we plot the estimated 95% credible interval for the over-parametrized substitution model. If over-parametrization has no impact, then all 95% credible interval sizes would follow the diagonal line (dashed line). The first row corresponds to the simulated trees with 16 taxa, while the second row corresponds to the simulated trees with 64 taxa. The mean branch lengths (BL) for the data sets are on top of each column. The two first columns display the data sets with 100 sites, the two other columns show the data sets with 1000 taxa. The symbols represent the different prior settings for the inference under GTR+Γ+I (Table 2). Separate plots for each model are shown in Figures S2-5.

We noticed the largest deviation between the true model (JC) and the over-parameterized models (GTR+Γ+I) when we used a mean of 0.02 for the branch lengths in our simulated trees. The estimated 95% credible interval was larger for the over-parameterized models. The trees simulated with a branch length prior with mean 0.02 had shorter branches and the prior distribution on branch lengths was further away from the true values (Table 1 and 2, Figure 6 and 7). The trees simulated with a branch length prior with mean 0.1 had a matching prior distribution in the inference. Thus, we observed some interaction between the branch length prior and the prior distribution on the substitution model parameters.

**Figure 7:**
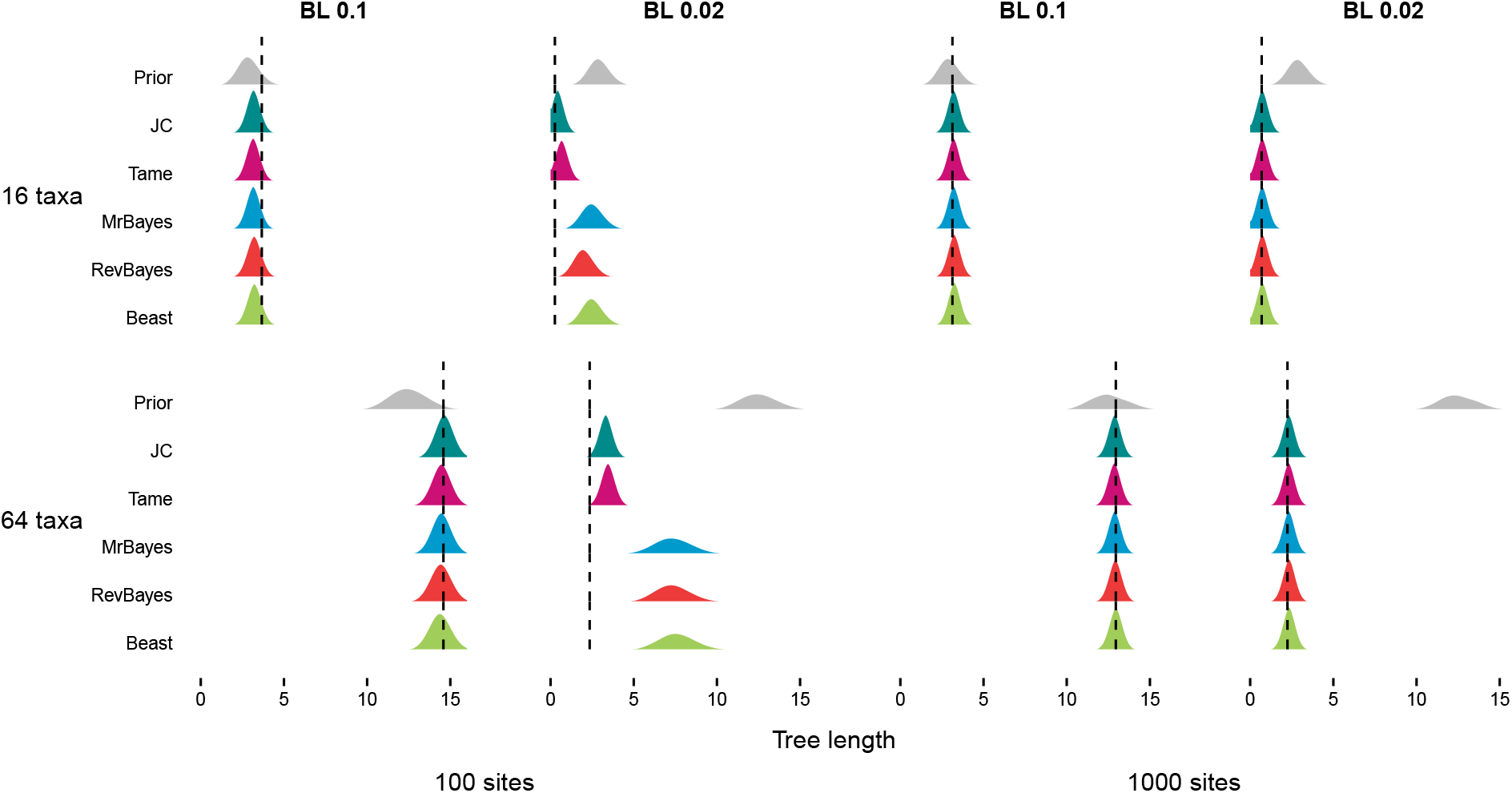
Comparison between prior and posterior distributions for the tree length. The first row corresponds to a simulated tree with 16 taxa, while the second row corresponds to a simulated tree with 64 taxa. The mean branch lengths (BL) for the data sets are on top of each column. The two first columns display the data sets with 100 sites, the two other columns show the data sets with 1000 taxa. The posterior distributions represented correspond to the inference performed under the true model (JC) and the over-parameterized model (GTR+Γ +I) with the 4 different prior settings (Table 2).

We showed one example data set for each simulated scenario to demonstrate more clearly the impact of substitution model over-parameterization on the estimated tree length (Figure 7). We observed that the posterior distribution inferred under the GTR+Γ+I substitution model with the `Tame` prior settings matches the posterior distribution inferred under the JC substitution model (the true model) for all scenarios. We observed the most extreme deviation between the posterior distribution inferred under the JC substitution model (the true model) and the inferred posterior distribution of the tree length under the `MrBayes`, `RevBayes` and `BEAST2` prior scheme for 64 taxa, 100 sites and mean branch length 0.02. In this scenario, the posterior distribution of the over-parameterized substitution model is widest and shifted in location. Thus, for the `MrBayes`, `RevBayes` and `BEAST2` prior schemes we observed biases and more uncertainty in the estimated parameters.

We plotted the estimated posterior distribution of the rate categories for the among site rate variation model to elucidate the problem of overestimated posterior distribution of the tree length and interaction of parameters (Figure 8). Note that under the true model, all four rate categories should be equal to 1.0.

**Figure 8:**
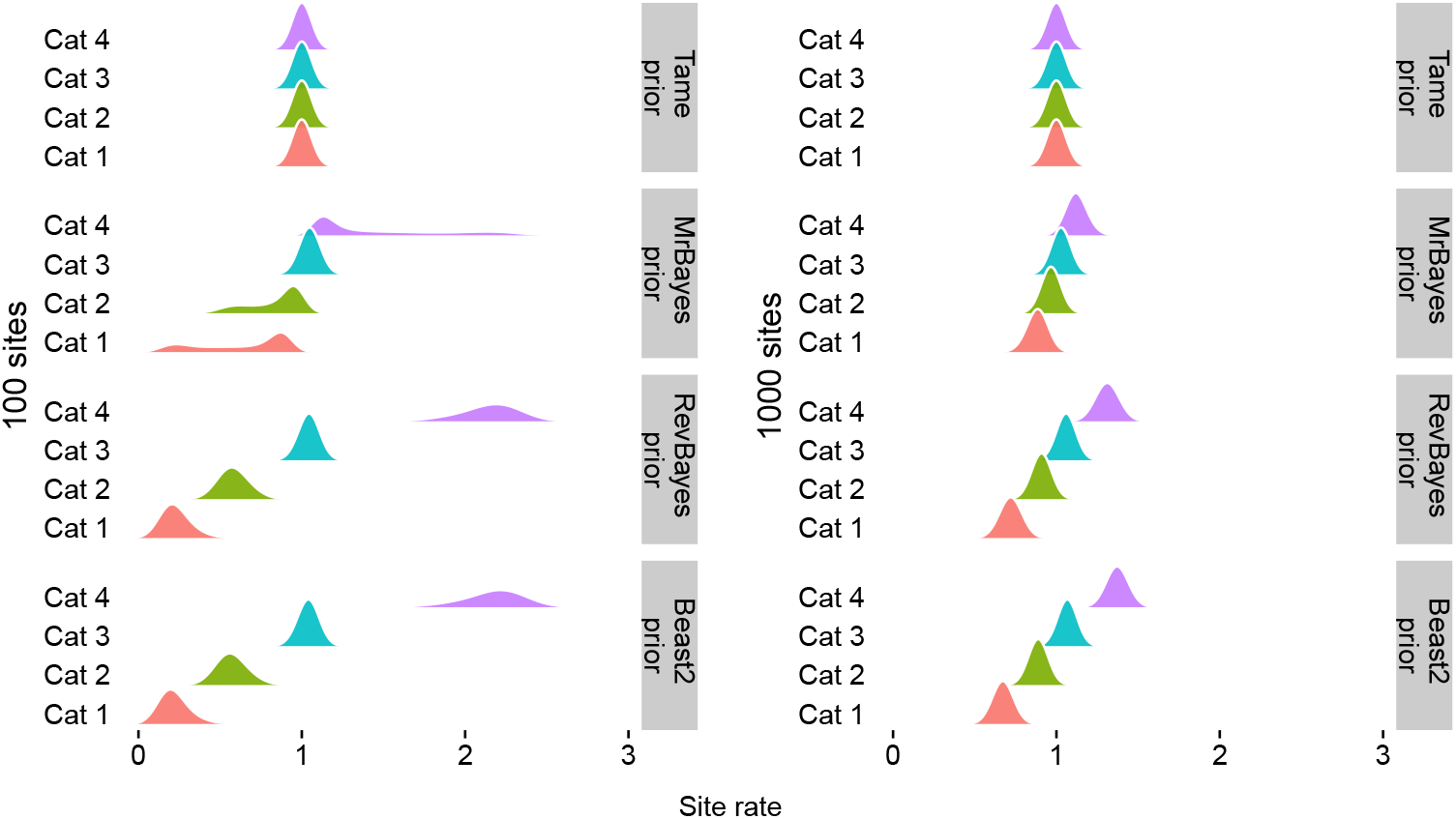
Posterior distributions for the rate categories for the among site rate variation model. The posterior distributions correspond to the inference performed under the over-parameterized model (GTR+Γ +I) with the four different prior settings (Table 2). The left panel shows the posterior distributions for one replicate data with 64 taxa, 100 sites and mean branch length 0.02, whereas the right panel the replicate data has 1000 sites. The posterior distributions are more sensitive to the prior (Figure 3) when the dataset was smaller.

The `MrBayes`, `RevBayes` and `BEAST2` prior settings yielded four gamma quartiles with slight (`MrBayes`) to large (`RevBayes` and `BEAST2`) differences *a posteriori* in relative site rates (Figure 8). When the sites are estimated to fall into the lower rate category, then longer branches are required to obtain the same amount substitutions. This results in larger branch lengths, as seen in Figures 6 and 7.

Additionally, we tested all possible combinations of the JC and GTR substitution models with the ASRV model and/or the invariant sites model to further evaluate the effect of the inappropriate prior distribution on *α* (Figures S8-S11). For each simulation setting (Table 1) we randomly selected one example simulated dataset and then performed the inference under the following models: JC+I, JC+Γ, JC+Γ+I, GTR, GTR+I, GTR+Γ, GTR+Γ+I. The results show that the 95% credible interval for the tree length was overestimated only when we added the ASRV model with the `MrBayes`, `RevBayes` and `BEAST2` prior settings. This result further corroborates that the bias on the tree length is exclusively linked with the prior distribution on *α* of the ASRV model.

Finally, we observed no biases in the accuracy of the tree length (Figure 9) for the inference under the partition model. Recall that we used the `Tame` prior settings for each data subset in our partition model, and used only datasets with 1000 sites equally divided into two subsets of each 500 sites. Since our results using the `Tame` prior on very small datasets of 100 sites showed no impact of substitution model over-parameterization (Figure 9), it is expected that the over-splitted and over-parameterized substitution model produces robust estimates of the tree length. In conclusion, Figures 6 and 9 demonstrate that Bayesian inference of phylogeny is robust to substitution and partition model over-parameterization; however, only if well behaved prior distributions are chosen.

**Figure 9:**
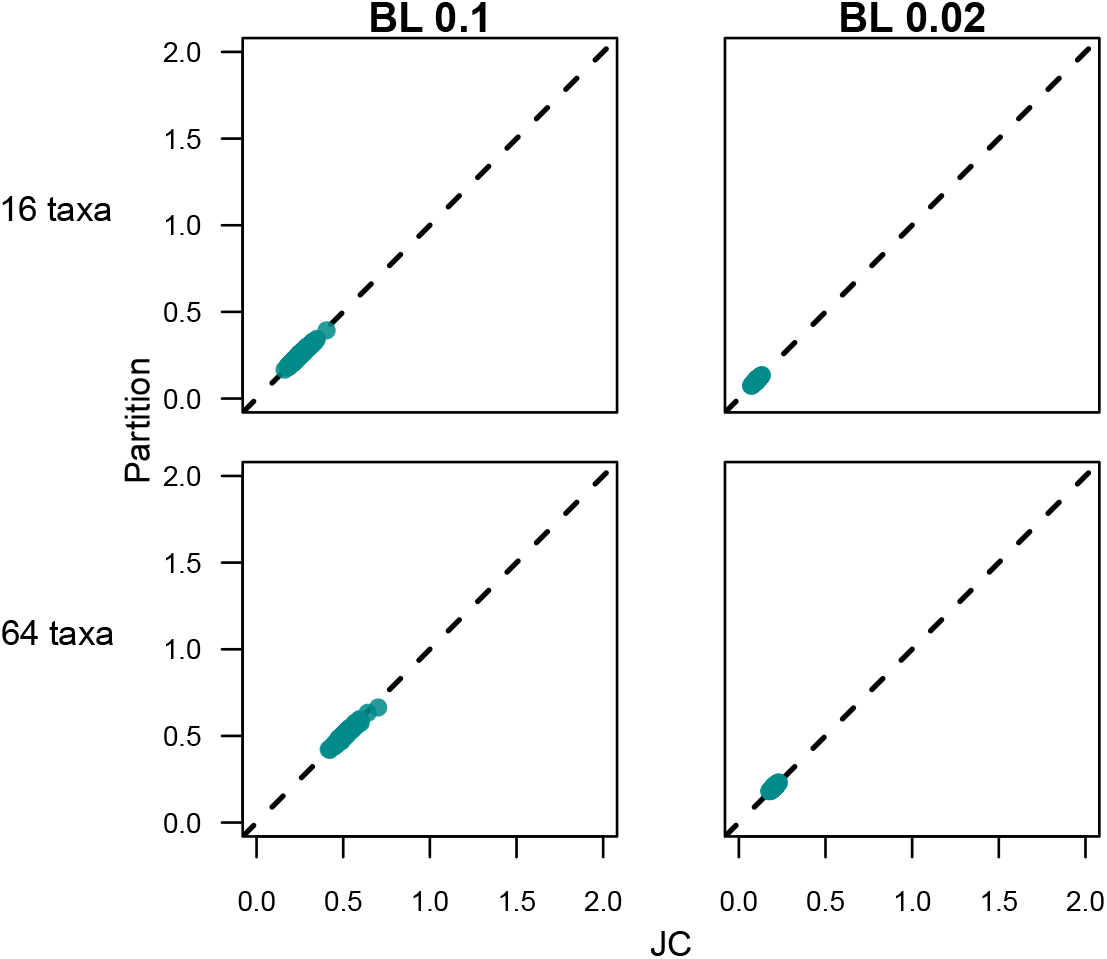
The width of the 95% credible interval for the tree length for the inference with Jukes-Cantor (JC) against the inference with the over-splitted GTR+Γ+I partitioned model. The first row corresponds to the simulated trees with 16 taxa, while the second row corresponds to the simulated trees with 64 taxa. The mean branch lengths (BL) for the data sets are on top of each column. For the over-partitioned model, we only used data sets with 1000 sites because splitting 100 sites results into unrealistically small data subsets. On the x-axis we show the estimated 95% credible interval size for the JC substitution model (true model). On the y-axis we plot the estimated 95% credible interval for the over-partioned model. If over-parametrization has no impact, then all 95% credible interval sizes would follow the diagonal line (dashed line).

## 4 Discussion and Conclusions

The main purpose of selecting a substitution model is to avoid model misspecification. Assuming an under-parameterized substitution model has been shown to bias phylogenetic inference (Sullivan and Joyce 2005). However, the question whether over-parameterization of substitution models biases phylogenetic inferences has received much less attention. Under- and over-parameterization of substitution models could be avoided if we would know the true substitution model. Since we do not know the true substitution model, it is common practice to perform substitution model selection before estimating a phylogeny. Common substitution model selection approaches, *e.g*., `ModelTest` (Posada and Crandall 1998), `jModelTest` (Posada 2008) and `jModelTest 2` (Darriba et al. 2012), employ shortcuts (*e.g*., do not take uncertainty in parameter estimate into account and optimize only some parameters) and are philosophically questionable (*e.g*., they require a phylogeny to perform the model selection step). In this manuscript we argue that substitution model selection is not necessary and can be avoided if the most complex substitution model (*e.g*., the GTR+Γ+I substitution model) with well behaved prior distribution is applied.

Applying the most complex substitution model comes with the cost that both the likelihood calculation is slower and there are more parameters to be estimated. If we could be absolutely certain that a simpler substitution model suffices, then we could save computation time by applying this simpler substitution model. However, given the shortcuts and philosophical shortcomings of substitution model selection procedure and uncertainty/disagreement in selection substitution models (Abadi et al. 2019), we argue that it is safer to err on the side of a too complex substitution model at a small amount of extra computational cost. Similarly, we argue that model averaging approaches (*e.g*., Huelsenbeck et al. 2004; Bouckaert and Drummond 2017) are not necessary —because our results show that there is no difference in estimated phylogenies between different models if well behaved prior distributions are chosen— and only unnecessarily increase computational demands due to more complex model averaging algorithms (*e.g*., reversible jump MCMC which is prone to poor MCMC convergence).

Furthermore, multi-locus datasets are used in phylogenetics since several decades where each locus evolves under a substitution process that is either independent or shared among loci (Nylander et al. 2004). The shortcuts taken to select the best partition model are more severe due to computational reasons (*e.g*., Frandsen et al. 2015) and their impact is less explored. In our simulation study we explored an extreme scenario where both the substitution model was over-parameterized and the locus was over-splitted. In line with our results on substitution model over-parameterization of single loci, we found that Bayesian analysis with a well behaved prior distribution does not bias phylogenetic inference. That means, one does not need to worry about assuming too many division of the data into subsets, although more data subsets come with a higher computational cost owed to the additional parameters.

In general, our results show that over-parameterization is not a problem in Bayesian inference if well behaved prior distributions are chosen, and these results could apply to models beyond substitution models. On the contrary, standard phylogenetic models are likely to be over-simplified and inadequate (Doyle et al. 2015; Höhna et al. 2018; Richards et al. 2018). There exist several extensions to standard phylogenetic substitution models, such as the CAT model (Lartillot and Philippe 2004) and Markov modulated substitution models (Baele et al. 2021). None of these more complex models are contained in substitution model selection approaches and our efforts should go into developing and testing more realistic substitution models.

Several previous studies have shown that Bayesian phylogenetic inferences can produce unrealistic long trees (Brown et al. 2010; Marshall 2010; Rannala et al. 2012; Zhang et al. 2012). These studies identified the prior distribution on the tree length as being responsible for the unrealistically large posterior estimates of the tree length. In our simulations, we noticed that if the true value for the tree length was far outside the center of prior distribution, then estimates of the tree length using phylogenetic analysis with the default prior settings for `MrBayes`, `BEAST2` and `RevBayes` were significantly biased. These previous studies by Brown et al. (2010), Marshall (2010) and Zhang et al. (2012) used `MrBayes` and therefore the results are likely influenced by an interaction of the tree length prior distribution and ASRV prior distribution (see also Ekman and Blaalid (2011)). Our proposed prior distribution (the `Tame` prior setting) does not show this interaction between tree length prior distribution and ASRV prior distribution and might alleviate the problem of unrealistically long trees.

In this manuscript we focused exclusively on over-parameterization of substitution models and the choice of prior distributions in a Bayesian inference framework. Our results may not be directly comparable to a Maximum likelihood (ML) framework and over-parameterization could still be a problem. In a ML framework, nuisance parameters, such as the substitution model parameters, are estimated to a single value that maximizes the likelihood instead of integrating over the uncertainty, which is done in a Bayesian framework. Therefore it is possible that a Bayesian inference is less impacted by over-parameterization as the uncertainty in the rate variation among sites could be large, but a ML inference on the other hand is impacted. A similar simulation study as ours could provide an answer to the question if over-parameterization is a problem for phylogenetic inference in a Maximum likelihood framework.

In conclusion, we found that over-parameterization is not a problem for Bayesian phylogenetic inference and does not bias tree topology estimates or branch length estimates if well behaved prior distributions are chosen. Our results corroborate previous findings by Huelsenbeck and Rannala (2004) and Lemmon and Moriarty (2004) on the robustness of estimating phylogenetic trees using over-parameterized substitution models. Here we additionally explored the impact of substitution model over-parameterization on estimates of the tree length (and thus by proxy on branch lengths) under different prior settings. We show that tree lengths estimates are more sensitive to substitution model over-parameterization and the choice of prior distributions. These prior distribution might result into unforeseen side-effects, for example, an informative prior distribution on the among site rate variation leads to more sensitivity to the prior on the tree length. We propose and tested a new choice of prior distribution, the `Tame` priors, which are well behaved. Our new choice of prior distribution can be applied to partition models and renders selection of the best partition scheme unnecessary. In general, substitution models are most likely too simple and our worries should focus on developing more realistic substitution models instead of selecting between a set of unrealistic substitution models.

## 5 Acknowledgements

We would like to thank Ronja Billenstein, Killian Smith and Bjørn Kopperud for feedback on the manuscript. This research was supported by the Deutsche Forschungsgemeinschaft (DFG) Emmy Noether-Program HO 6201/1-1 awarded to SH.

