## Supplementary Material for "Bayesian inference of phylogeny is robust to substitution model over-parameterization"

#### Supporting Information

Luiza Guimarães Fabreti[1,2] and Sebastian Höhna[1,2,*]

[1] *GeoBio-Center, Ludwig-Maximilians-Universität München, Richard-Wagner Straße 10, 80333 Munich, Germany*
[2] *Department of Earth and Environmental Sciences, Paleontology & Geobiology, Ludwig-Maximilians-Universität München, Richard-Wagner Straße 10, 80333 Munich, Germany*

### Contents

**S1 Variable sites in simulated data sets** **3**

**S2 Credible interval for the Tree Length** **4**

**S3 Mean Tree Length for 100 sites and short branch lengths** **8**

**S4 Further exploration of model combinations** **10**

**S5 Convergence assessment example** **13**

### S1 Variable sites in simulated data sets

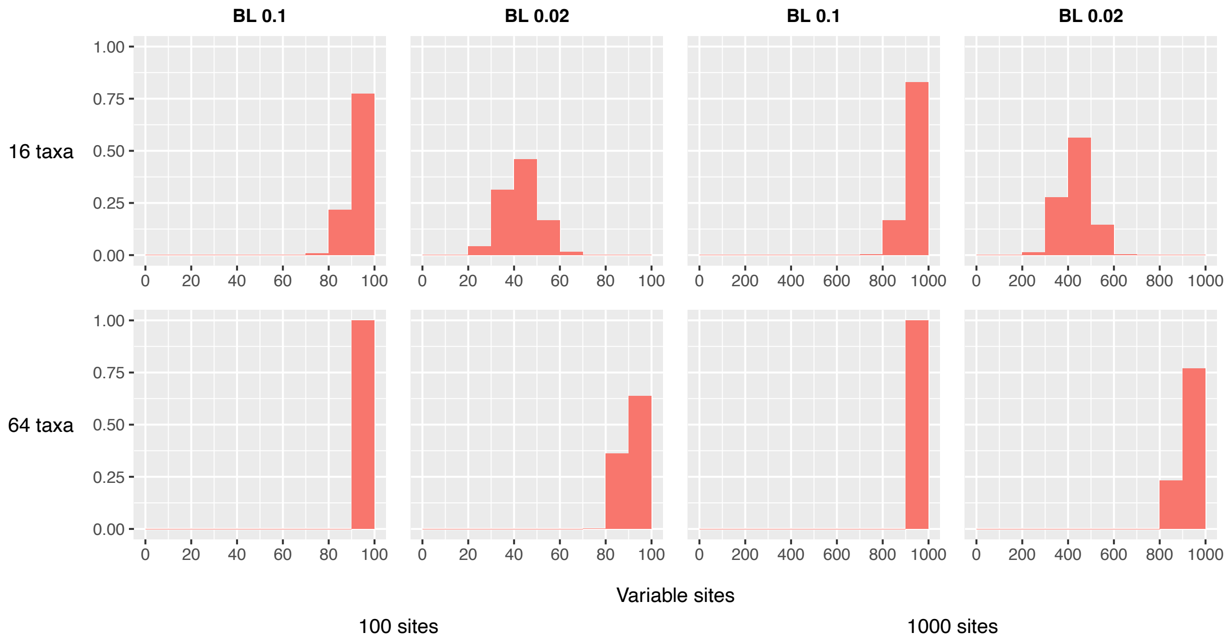


**Figure S1:** Histogram of relative number of variable sites in the data sets for each simulation scenario. The data were simulated in `RevBayes` [5]. The first row corresponds to the simulated trees with 16 taxa, while the second row corresponds to the simulated trees with 64 taxa. The mean branch lengths (BL) for the data sets are on top of each column. The two first columns display the data sets with 100 sites, the two other columns show the data sets with 1000 taxa.

#### S2 Credible interval for the Tree Length

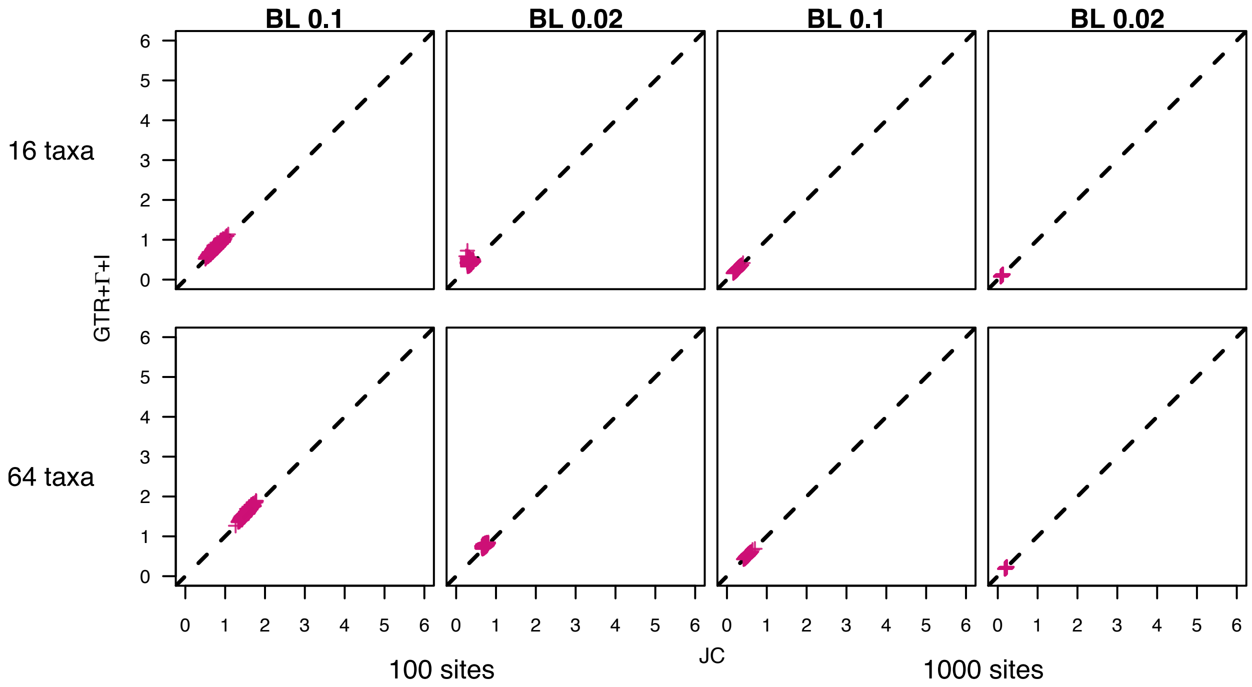


**Figure S2:** The 95% credible interval for the tree length for the inference with Jukes-Cantor [JC, 6] against the inference with GTR+Γ+I [8, 9, 3] under the `Tame` prior. On the x-axis we show the estimated 95% credible interval size for the JC substitution model (true model). On the y-axis we plot the estimated 95% credible interval for the over-parametrized substitution model. The first row corresponds to the simulated trees with 16 taxa, while the second row corresponds to the simulated trees with 64 taxa. The mean branch lengths (BL) for the data sets are on top of each column. The two first columns display the data sets with 100 sites, the two other columns show the data sets with 1000 taxa.

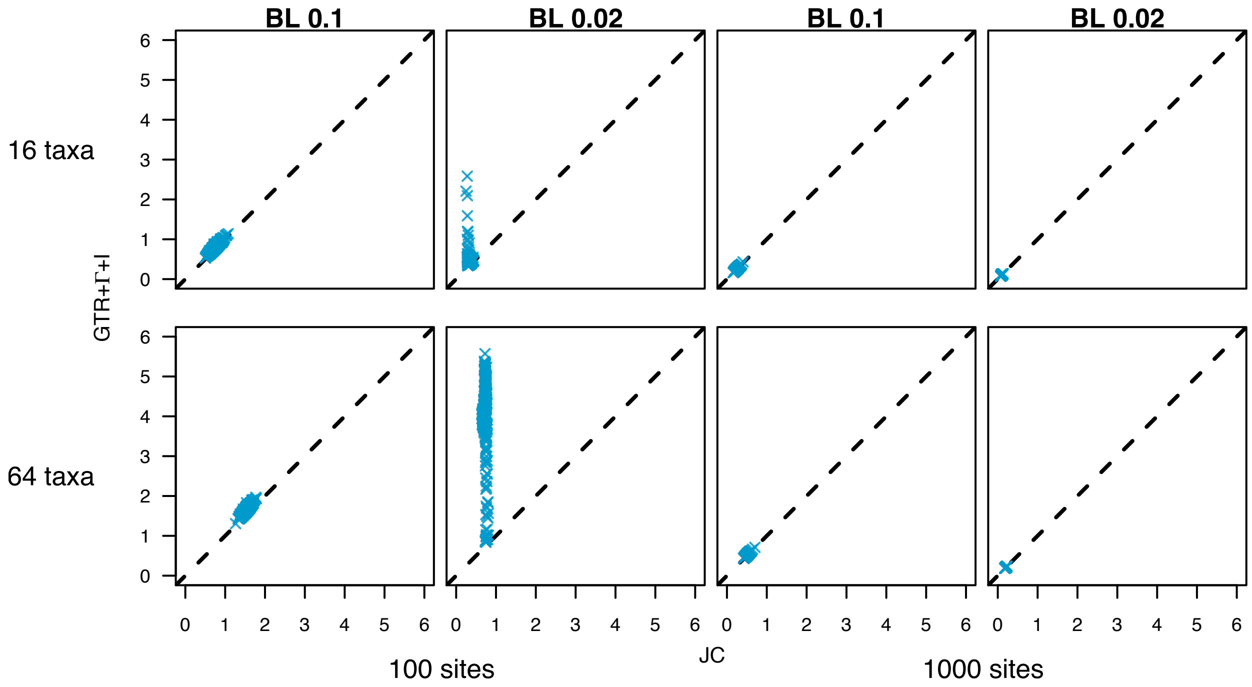


**Figure S3:** The 95% credible interval for the tree length for the inference with Jukes-Cantor [JC, 6] against the inference with GTR+Γ+I [8, 9, 3] under the `MrBayes` prior [7]. On the x-axis we show the estimated 95% credible interval size for the JC substitution model (true model). On the y-axis we plot the estimated 95% credible interval for the over-parametrized substitution model. The first row corresponds to the simulated trees with 16 taxa, while the second row corresponds to the simulated trees with 64 taxa. The mean branch lengths (BL) for the data sets are on top of each column. The two first columns display the data sets with 100 sites, the two other columns show the data sets with 1000 taxa.

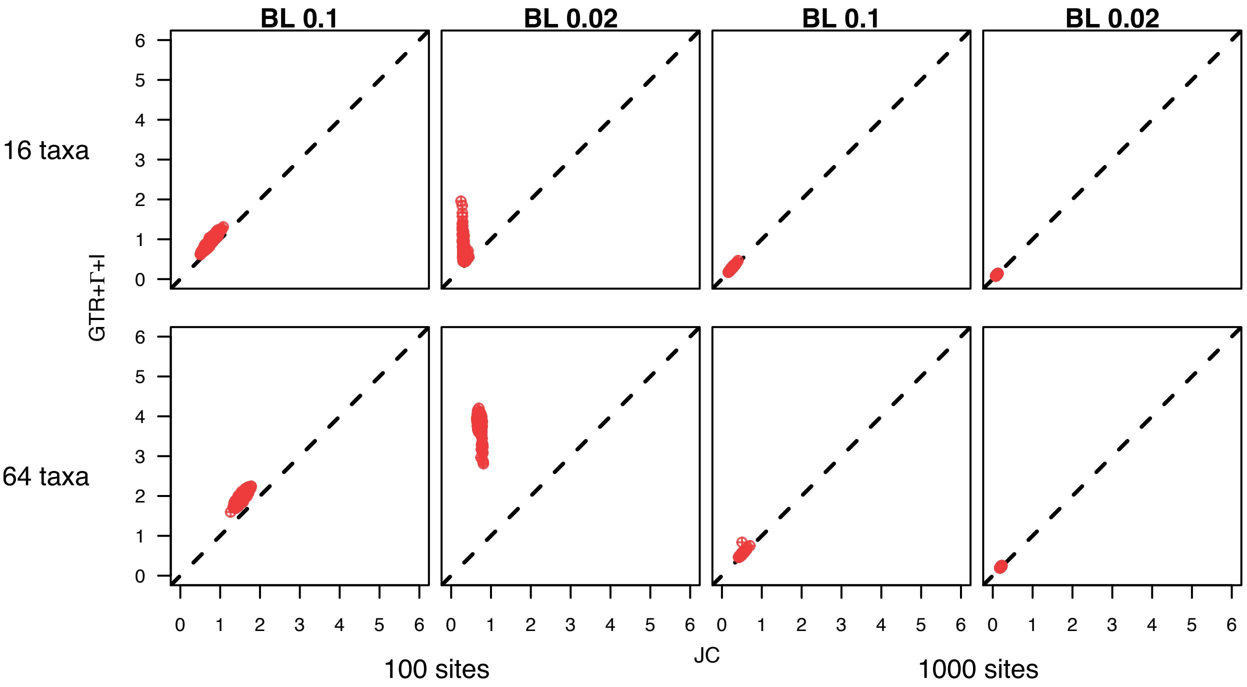


**Figure S4:** The 95% credible interval for the tree length for the inference with Jukes-Cantor [JC, 6] against the inference with GTR+Γ+I [8, 9, 3] under the `RevBayes` prior [4]. On the x-axis we show the estimated 95% credible interval size for the JC substitution model (true model). On the y-axis we plot the estimated 95% credible interval for the over-parametrized substitution model. The first row corresponds to the simulated trees with 16 taxa, while the second row corresponds to the simulated trees with 64 taxa. The mean branch lengths (BL) for the data sets are on top of each column. The two first columns display the data sets with 100 sites, the two other columns show the data sets with 1000 taxa.

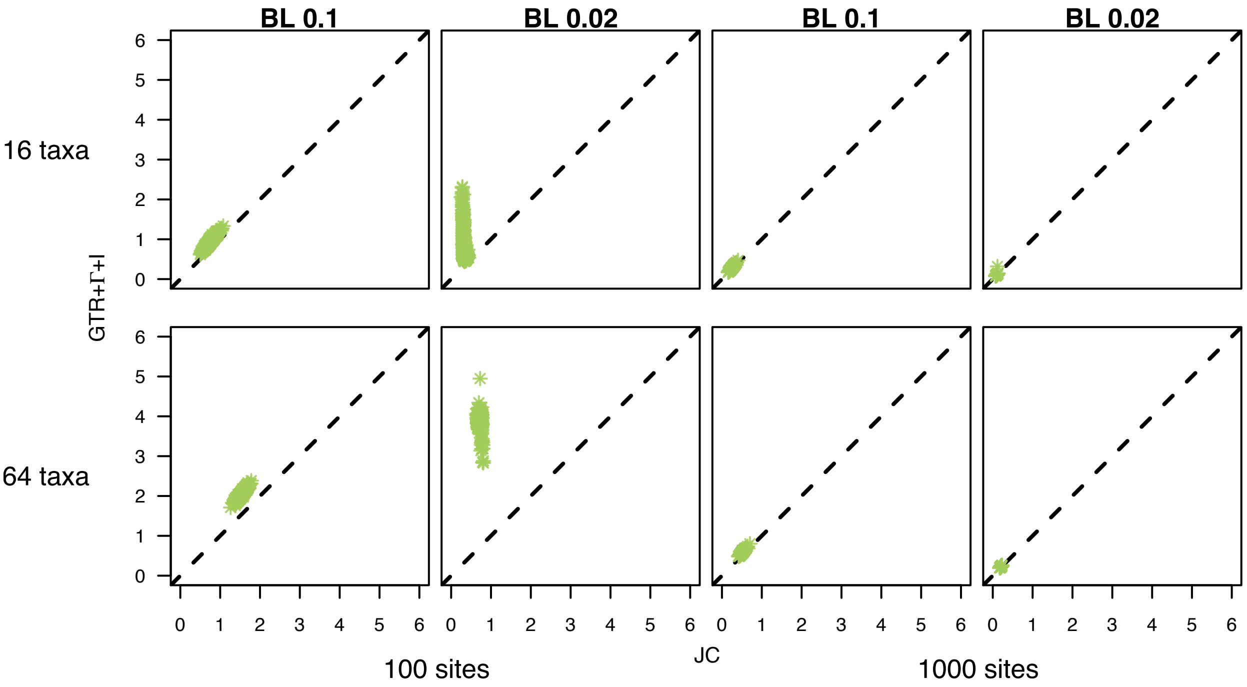


**Figure S5:** The 95% credible interval for the tree length for the inference with Jukes-Cantor [JC, 6] against the inference with GTR+Γ+I [8, 9, 3] under the BEAST2 prior [1]. On the x-axis we show the estimated 95% credible interval size for the JC substitution model (true model). On the y-axis we plot the estimated 95% credible interval for the over-parametrized substitution model. The first row corresponds to the simulated trees with 16 taxa, while the second row corresponds to the simulated trees with 64 taxa. The mean branch lengths (BL) for the data sets are on top of each column. The two first columns display the data sets with 100 sites, the two other columns show the data sets with 1000 taxa.

#### S3 Mean Tree Length for 100 sites and short branch lengths

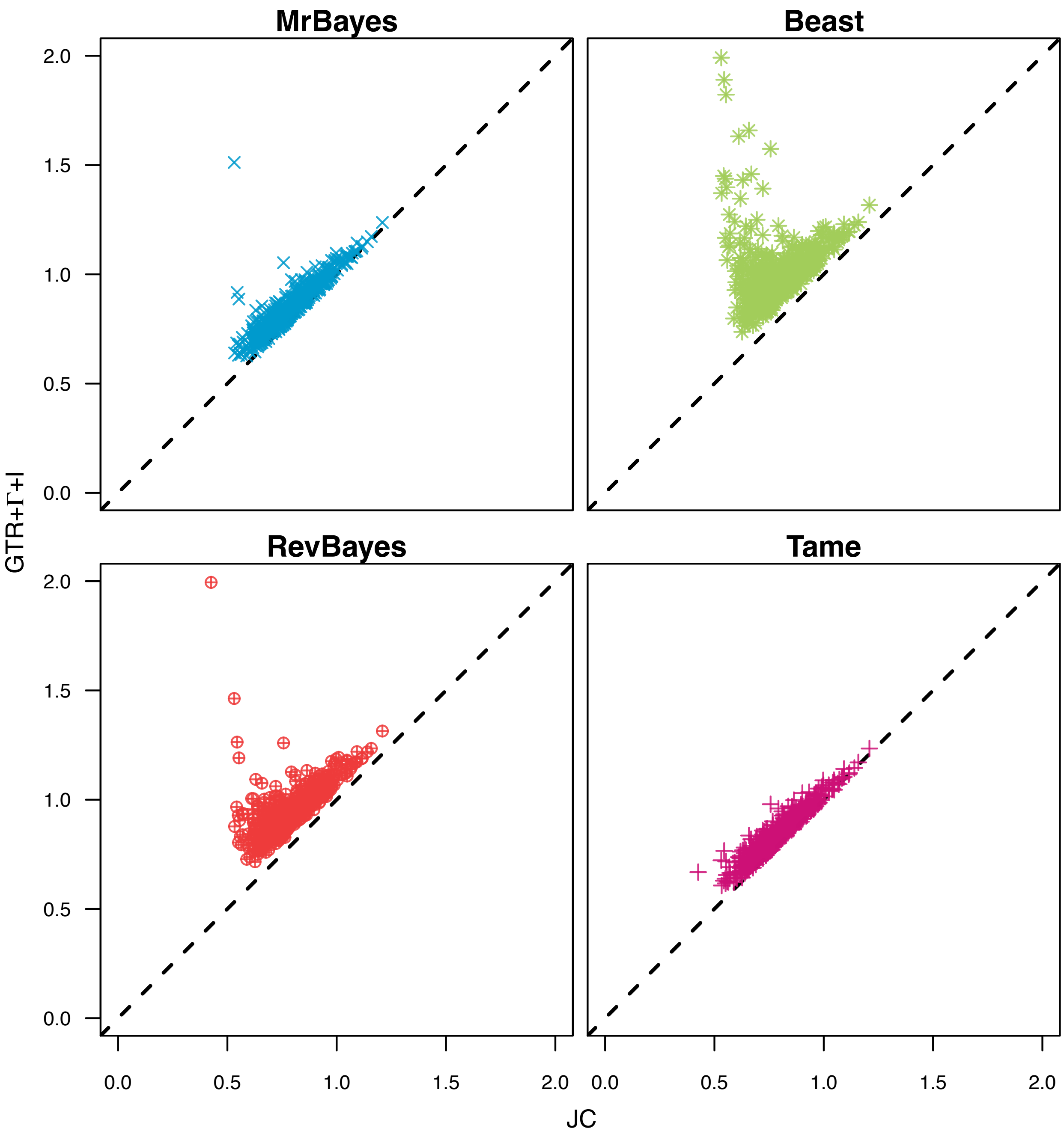


**Figure S6:** Comparison of mean tree length between [JC, 6] and GTR+Γ+I [8, 9, 3] for the data sets with 16 taxa, 100 sites and mean branch length 0.02. The x-axis represents the mean tree length for the inference under JC, while the y-axis represents the mean tree length for the inference under GTR+Γ+I. Each plot displays the means for the inference with the four different prior schemes.

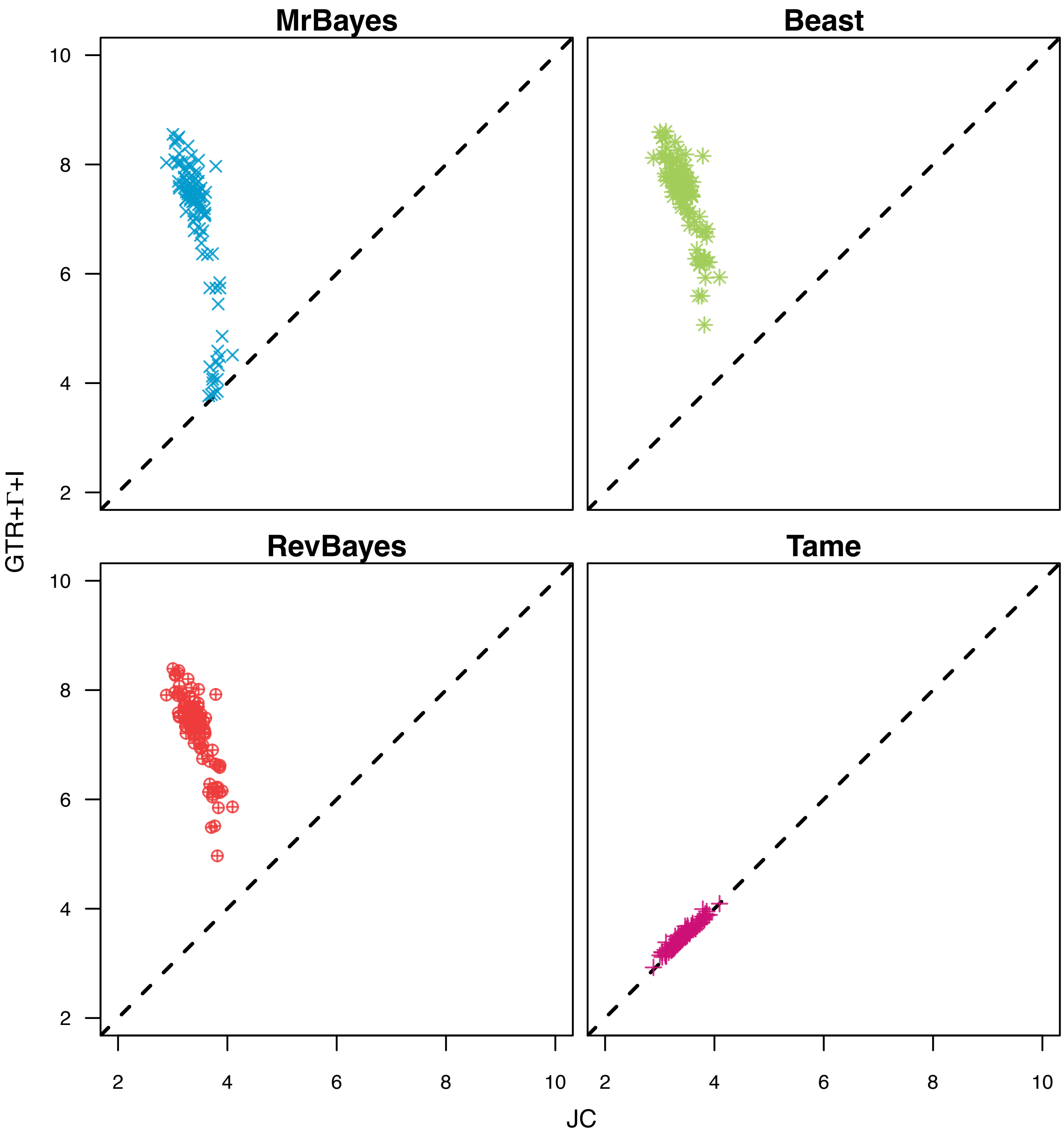


**Figure S7:** Comparison of mean tree length between [JC, 6] and GTR+Γ+I [8, 9, 3] for the data sets with 64 taxa, 100 sites and mean branch length 0.02. The x-axis represents the mean tree length for the inference under JC, while the y-axis represents the mean tree length for the inference under GTR+Γ+I. Each plot displays the means for the inference with the four different prior schemes.

#### S4 Further exploration of model combinations

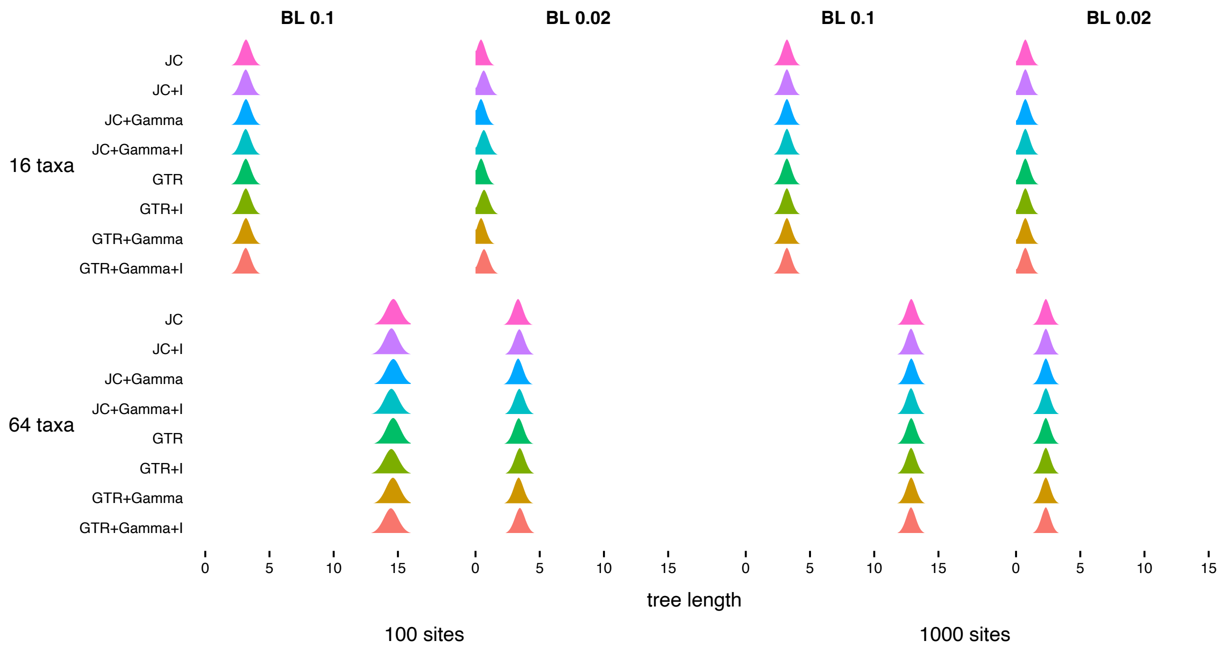


**Figure S8:** Posterior distributions for tree length for one example of each data set. Each line corresponds to a different model (JC, JC+I, JC+Γ, JC+Γ+I, GTR, GTR+I, GTR+Γ, GTR+Γ+I; [6, 8, 9, 3]). The GTR models followed the `Tame` prior setting. The first row corresponds to the simulated trees with 16 taxa, while the second row corresponds to the simulated trees with 64 taxa. The mean branch lengths (BL) for the data sets are on top of each column. The two first columns display the data sets with 100 sites, the two other columns show the data sets with 1000 taxa.

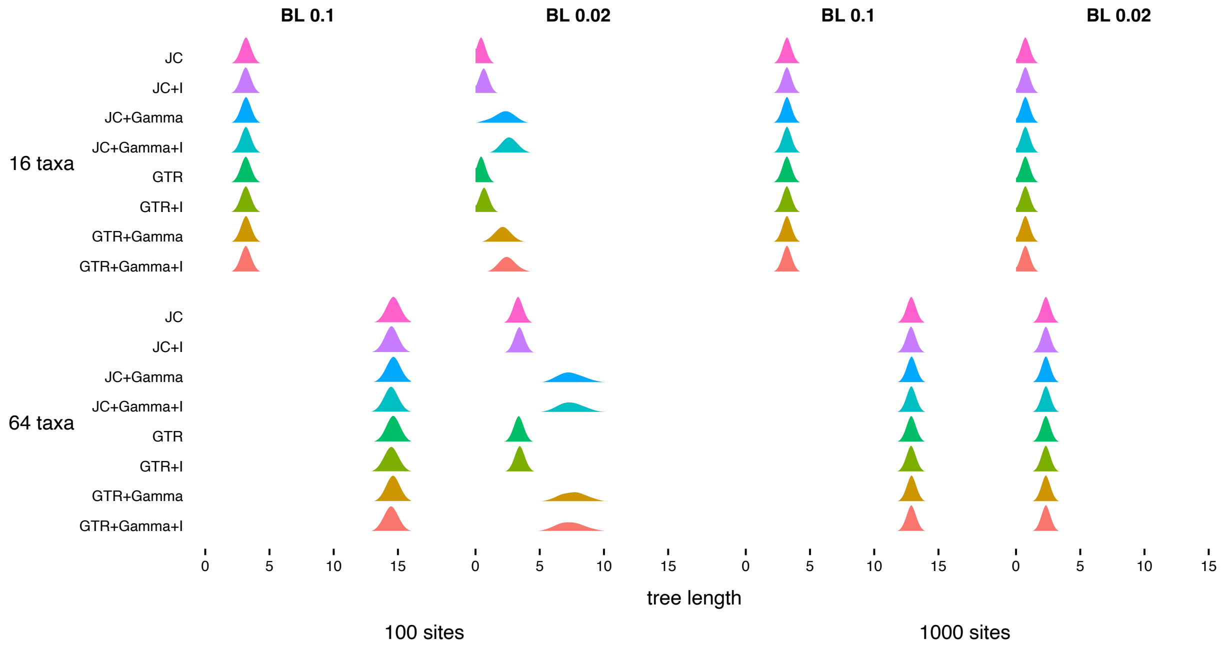


**Figure S9:** Posterior distributions for tree length for one example of each data set. Each line corresponds to a different model (JC, JC+I, JC+Γ, JC+Γ+I, GTR, GTR+I, GTR+Γ, GTR+Γ+I; [6, 8, 9, 3]). The GTR models followed the `MrBayes` prior setting. The first row corresponds to the simulated trees with 16 taxa, while the second row corresponds to the simulated trees with 64 taxa. The mean branch lengths (BL) for the data sets are on top of each column. The two first columns display the data sets with 100 sites, the two other columns show the data sets with 1000 taxa.

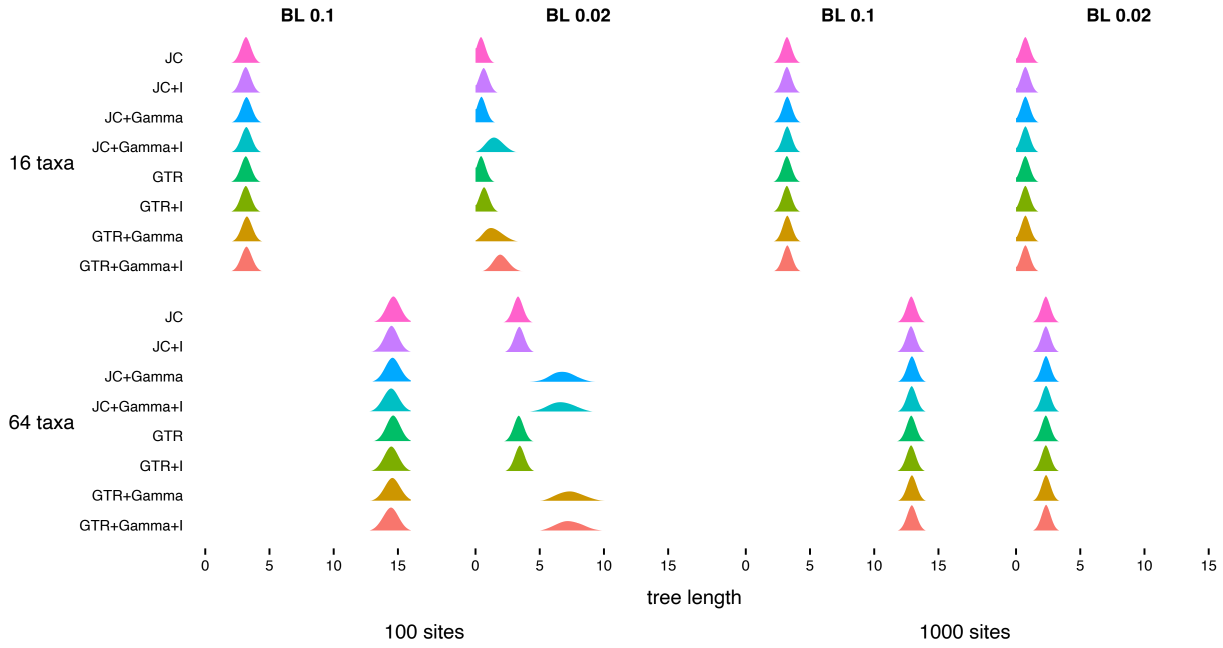


**Figure S10:** Posterior distributions for tree length for one example of each data set. Each line corresponds to a different model (JC, JC+I, JC+Γ, JC+Γ+I, GTR, GTR+I, GTR+Γ, GTR+Γ+I; [6, 8, 9, 3]). The GTR models followed the `RevBayes` [4] prior setting. The first row corresponds to the simulated trees with 16 taxa, while the second row corresponds to the simulated trees with 64 taxa. The mean branch lengths (BL) for the data sets are on top of each column. The two first columns display the data sets with 100 sites, the two other columns show the data sets with 1000 taxa.

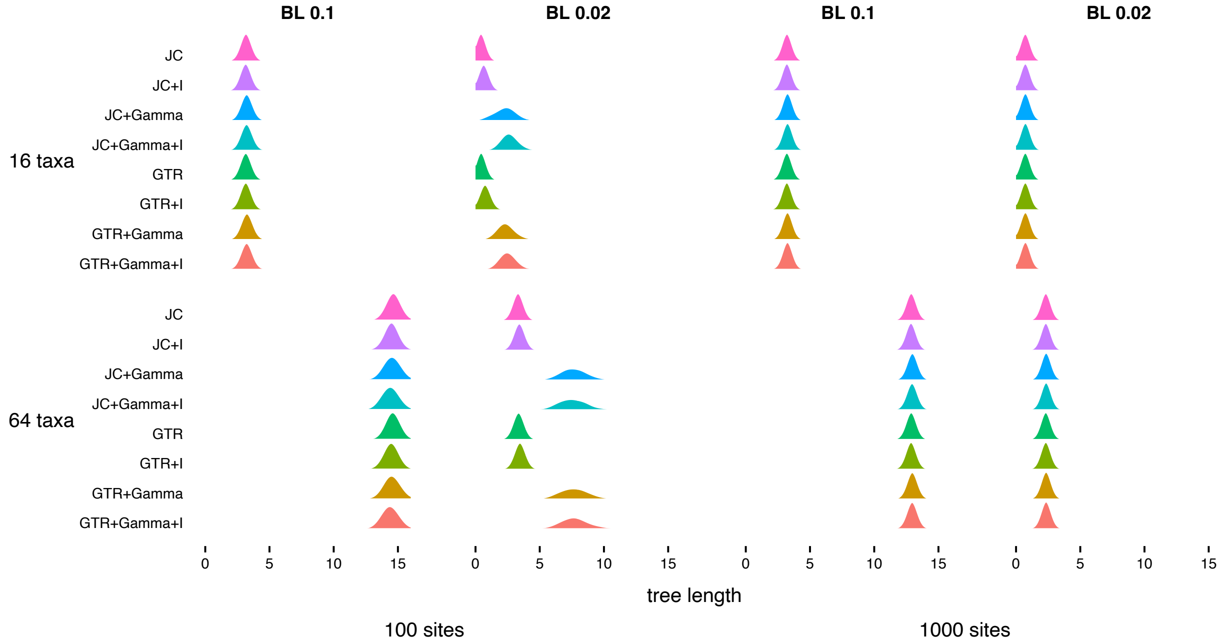


**Figure S11:** Posterior distributions for tree length for one example of each data set. Each line corresponds to a different model (JC, JC+I, JC+Γ, JC+Γ+I, GTR, GTR+I, GTR+Γ, GTR+Γ+I; [6, 8, 9, 3]). The GTR models followed the `BEAST2` [1] prior setting. The first row corresponds to the simulated trees with 16 taxa, while the second row corresponds to the simulated trees with 64 taxa. The mean branch lengths (BL) for the data sets are on top of each column. The two first columns display the data sets with 100 sites, the two other columns show the data sets with 1000 taxa.

### S5 Convergence assessment example

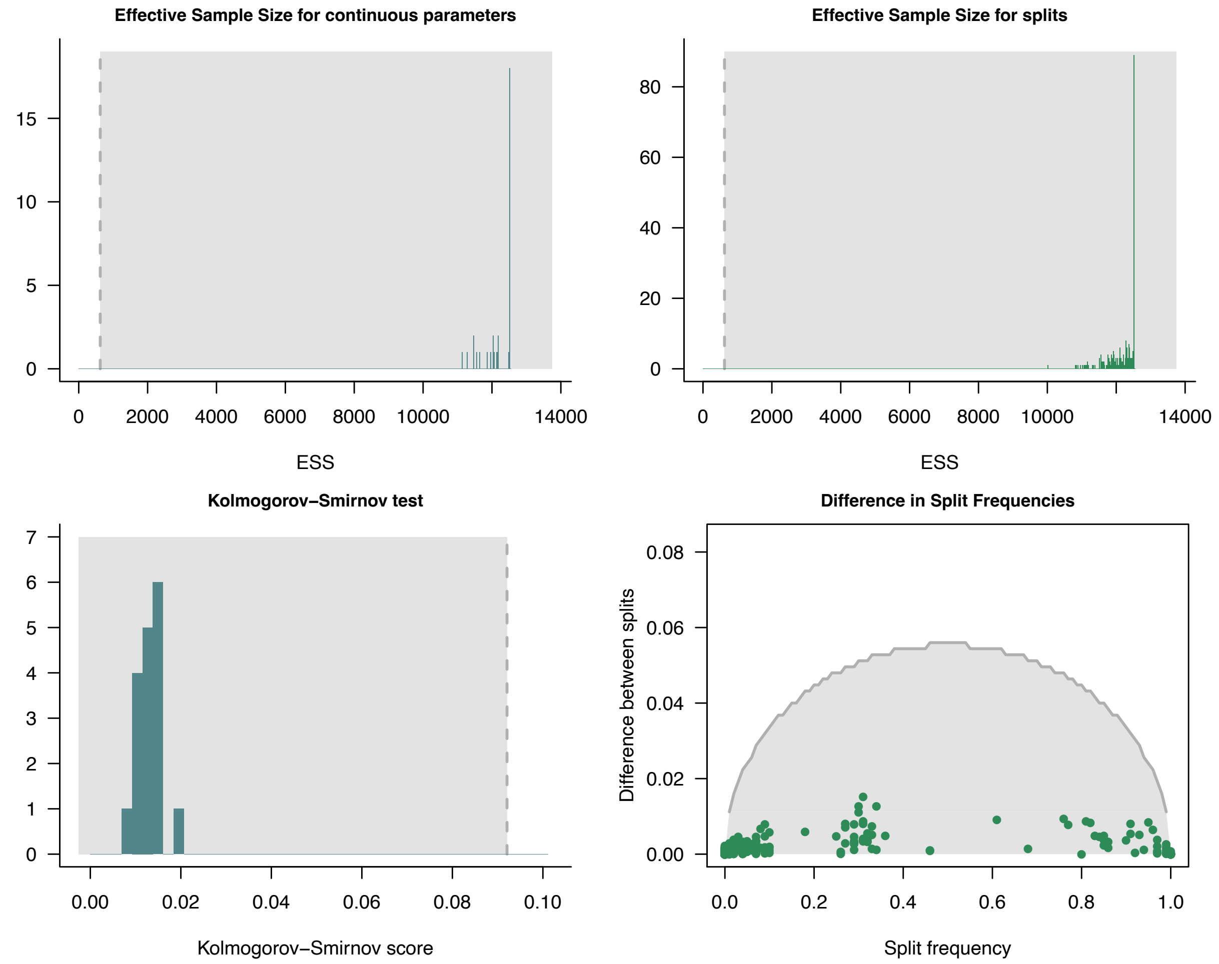


**Figure S12:** Convergence assessment plots using the package `Convenience` [2] for one example MCMC from Figure 7. The first column shows the plots for the assessment of convergence for the continuous parameters. The second column displays the assessment of the splits in the tree. In all plots the values are within the gray shaded area, which indicates that convergence was achieved.
